# Cell Communication Network factor 4 (CCN4/WISP1) shifts melanoma cells from a fragile proliferative state to a resilient metastatic state

**DOI:** 10.1101/487751

**Authors:** Wentao Deng, Audry Fernandez, Sarah L. McLaughlin, David J. Klinke

## Abstract

While deregulated intracellular signaling initiates melanoma, intercellular crosstalk within the tumor microenvironment, often coordinated by soluble factors, is essential for melanoma progression and metastasis. One such secreted matricellular protein, cellular communication network factor 4 (CCN4/WISP1), stimulates metastasis in other malignancies. Here, we report that CCN4 expression is associated progressively with reduced overall survival in patients with primary melanomas. To reveal the roles of CCN4 in melanoma progression, we used mouse melanoma models and knocked out *Ccn4* using a homology-directed repair CRISPR/CAS9 system to generate pools of *Ccn4*-knockout cells. *In vitro* assays supported previous findings using clones generated using a double nickase-based CRISPR/CAS9 system that CCN4 promoted an epithelial – mesenchymal-like transition in melanoma cells and stimulated invasion and metastasis. We also found that, while *Ccn4* knockout enhanced cell growth in optimal 2D culture conditions, the knockout suppressed certain cell survival signaling pathways and rendered cells less resistant to stress conditions. Tumor cell growth assays at sub-optimal conditions *in vitro*, quantitative analysis of tumor growth assays *in vivo*, and transcriptomics analysis of human melanoma cell lines suggested that CCN4 repressed cell growth and simultaneously enhanced cell survival. The collective role of CCN4 suggests a potential therapeutic target for limiting metastatic invasion in melanoma and a biomarker for metastatic potential.

## INTRODUCTION

While invasive melanoma accounts for only 1% of all skin cancer cases, it results in the vast majority (72%) of skin cancer deaths (1). Patient survival depends on the clinical stage at diagnosis, with 5-year survival rate of 98% for localized melanoma but 62% and 18% for regional and distant stage melanoma, respectively (1). Unfortunately, melanoma tends to disseminate early during tumor development (2)(3). The early dissemination and metastasis of melanoma results from a reversible switch between differentiated and invasive phenotypes (4)(5)(6). Complementing oncogenic signals, signals from the melanoma microenvironment play a key role, as they stimulate this phenotypic switch (5)(6)(7). Among the microenvironmental signals likely involved is <u>C</u>ellular <u>C</u>ommunication <u>N</u>etwork factor 4 (CCN4) (8–10)(11). Up-regulated by β-catenin transcriptionally, CCN4 is a cysteine-rich matricellular protein that is secreted by tumor cells and fibroblasts (8)(12). CCN4 promotes tumor cell proliferation, survival, migration/invasion and metastasis in a variety of human tumors such as breast, pancreatic, prostate, lung, colorectal, and brain cancers (13)(14).

In the context of melanoma, reports on CCN4 suggest a different role. For instance, Hashimoto et al. overexpressed CCN4 driven by a CMV promoter to conclude that CCN4 represses the *in vivo* growth and metastasis of a highly metastatic mouse melanoma line (8). In addition, Shao et al., using 1205Lu melanoma cells together with “Notch-engineered” fibroblasts, show that recombinant CCN4 inhibits melanoma growth *in vitro* and CCN4 secretion by adjacent fibroblasts represses melanoma growth (10). In contrast, our work using double nickase-based CRISPR/Cas9 systems to modify mouse and human melanoma cells demonstrate that CCN4 stimulates invasion and metastasis by promoting an Epithelial - Mesenchymal Transition (EMT)-like process (11). As double nickase-based CRISPR/Cas9 systems may introduce phenotypic bias, the objective of this study was to clarify the role of CCN4 in the context of melanoma using a homology directed repair-based CRISPR/Cas9 approach.

## RESULTS

### CCN4 is associated with reduced overall survival in primary melanoma patients

To assess the clinical implications of *CCN4* overexpression, we tested for possible connection between *CCN4* mRNA expression and survival of patients in skin cutaneous melanoma (SKCM) arm of the Cancer Genome Atlas (TCGA). Given the disconnect between the diagnosis and sample collection times among the SKCM samples, we used data from samples obtained at diagnosis from patients with primary melanoma and complete survival histories for statistical analysis. To explore an inverse relationship between *CCN4* expression and overall survival, a total of 95 primary samples with RNA-seq results were separated into four quartiles based on *CCN4* mRNA expression levels from low to high and analyzed using Kaplan-Meier survival curves (Fig.1). A Cox proportional hazards model estimated overall survival of these 95 patients with quartile of *CCN4* expression, tumor stage, and gender as covariates. Among CCN4 quartiles, there were no significant differences in either tumor stage or gender (p-value = 0.288). Using the p-values as a metric for the strength of evidence, the highest *CCN4* expression group was the only group with a likely increased hazard ratio over random chance (p-value = 0.084) compared with patients having the lowest *CCN4* expression, with tumor stage and gender having no significant prognostic value (tumor stage p-value > 0.99; gender p-value = 0.939). The hazard ratios were also progressively increased in each quartile linked with an increase in *CCN4* expression (Q1 HR: 1.00; Q2 HR: 1.15, 95% CI: 0.254 – 5.243, p-value = 0.852; Q3 HR: 1.96, 95% CI: 0.484 – 7.957, p-value = 0.345; Q4 HR: 3.14, 95% CI: 0.859 – 11.459, p-value = 0.084). Collectively, these results suggested that *CCN4* expression was progressively associated with a worse outcome in patients diagnosed with melanoma.

**FIGURE 1.**
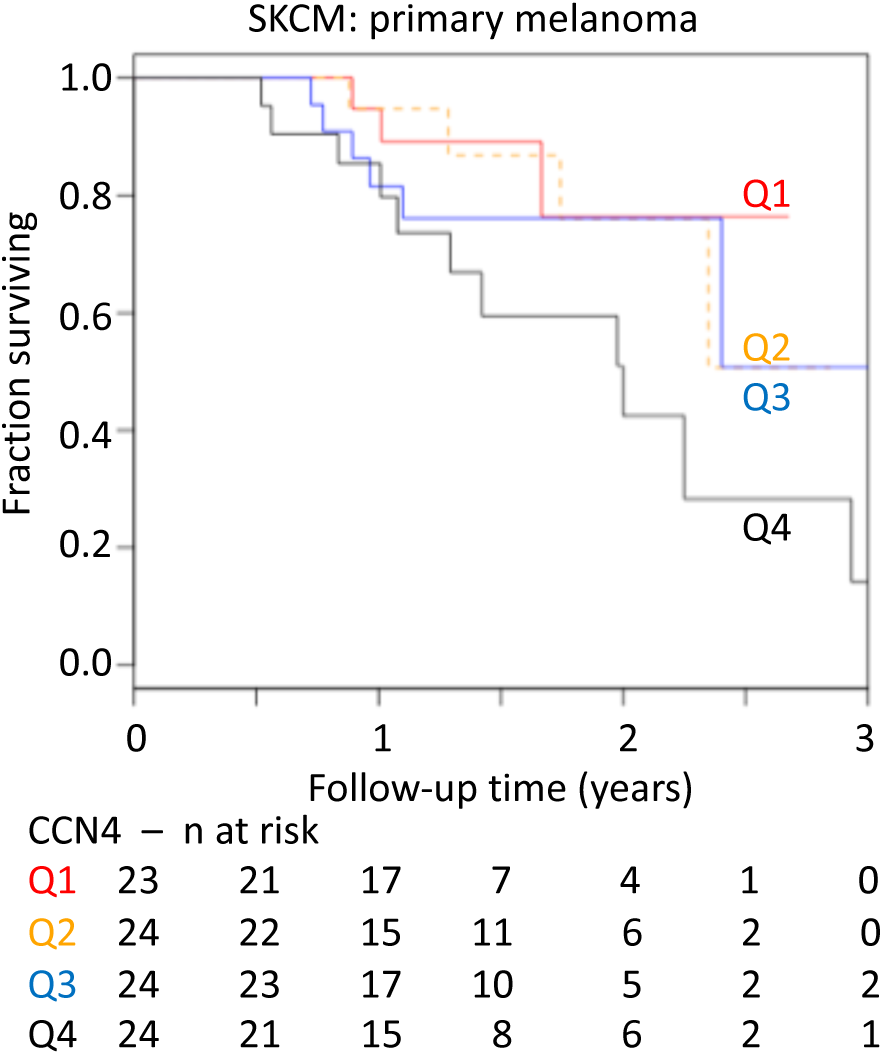
Kaplan-Meier estimates of overall survival for primary melanoma patients with increased *CCN4* expression. Data were from SKCM TCGA and patients with primary melanomas were separated into four quartiles (q1-q4) based on *CCN4* mRNA expression levels (low to high).

### Creation of *Ccn4*-Knockout Melanoma B16F0 and B16F10 Cell Pools

Previously, we observed that CCN4 stimulates melanoma cell invasion and metastasis *in vitro* and *in vivo* using mouse B16F10, YUMM1.7 and human RPMI-7951 melanoma lines (11). To investigate the role of CCN4 through loss-of-function strategy, we used a double nickase-based CRISPR/Cas9 system to target the *CCN4* gene at two different locations in each cell line. Although the results were generally consistent (11), off-target mutations caused by the DN-CRISPR system (15,16) remains as a potential source of bias, as the individual cell clones studied may contain unintended genetic backgrounds that do not exist in parental cells. In addition, the tendency for picking fast growing clones in the process of creating cell variants may also introduce a phenotypic bias in the knockout cell lines and limit generalization.

To overcome these potential sources of bias, we generated pools of mixed *Ccn4*-knockout cells from both B16F0 and B16F10 cells using a different CRISPR/Cas9 approach based on homology-directed repair (HDR). The HDR-CRISPR/Cas9 system contains knockout plasmids targeting three different sites in the genome and their corresponding HDR plasmids, which uses homology arm DNA to insert a puromycin-resistance gene after the site-specific double strand break (DSB) in the genomic DNA. Two 800 bp homology arms for each CRISPR targeting site and puromycin selection ensured creating specific and stable knockout cell pools (-KO). The mixture of knockout cells are expected to average out the off-target background and eliminate the potential proliferation bias that may have been present in the previous selection. The resulting (-KO) pools of modified cells contain one or more LoxP-flanked puromycin-resistance genes at its genomic *Ccn4* targeting sites, and the inserted cassettes were removed by Cre recombinase to generate (-KO’) cell pools. ELISA showed that the knockout system efficiently disrupted *Ccn4* expression in both B16F0 and B16F10 cells, and CCN4 secreted by knockout cells was only 0.09-7.8% of the original concentration (Fig.2A).

**FIGURE 2.**
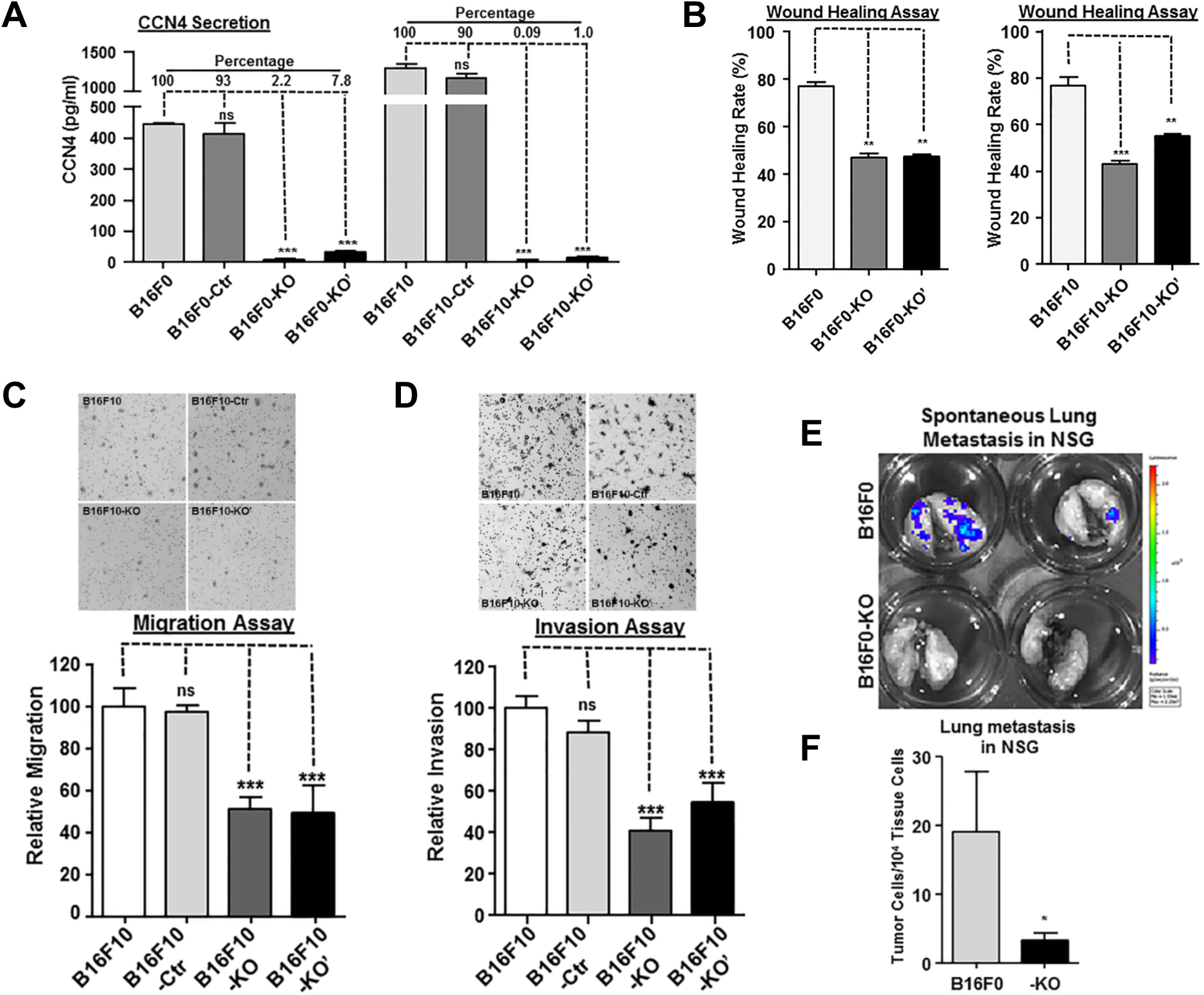
*Ccn4* knockout in mouse B16 melanoma cells represses wound healing, migration, invasion *in vitro* and tumor metastasis *in vivo*. **A.** CCN4 secretion after CRISPR/Cas9 knockout in mouse B16F0 and B16F10 melanoma cells. Cell culture medium were tested by ELISA after cells were plated for 48 hours. **B.** Wound healing of B16F0, B16F10 and their corresponding knockout cells. Scratches were created on 6-well plates in triplicate and the healing rate was calculated after 24 hours. **C.** Migration comparison of B16F10 and its knockout cells. Transwell migration assays were performed in triplicates and membranes were stained and counted after 24 hours. A representative migration staining from each group are shown on top. **D.** Invasion comparison of B16F10 and its knockout cells. Transwell invasion assays were performed in triplicates and membranes were stained and counted after 24 hours. A representative invasion staining from each group are shown on top. **E.** *In vivo* spontaneous lung metastasis assay with *ex vivo* bioluminescence imaging. NOD-SCID IL2Rgamma^null^ (NSG) mice with subcutaneous injection of B16F0 or B16F0-KO cells were terminated at day 21 post injection (n=3 in each group) for the imaging. Representative results were shown. **F.** *In vivo* spontaneous lung metastasis assay via real time genomic qPCR. Lung samples described in panel E were quantitatively analyzed and tumor metastatic burdens were presented as tumor cell number within 10,000 mouse tissue cells. Statistical significance was determined by unpaired two-tailed Student’s t-test, where a p-value < 0.05 was considered significant and asterisks was used to indicate calculated range in p-values. *: p-value < 0.05; **: p-value < 0.01; ***: p-value < 0.001; and ns: not significant.

### *Ccn4* Knockout in B16 Cells Repressed Wound Healing, Migration, Invasion *in Vit*ro and Tumor Metastasis *in Vivo*

With the new *Ccn4*-knockout B16 cells, we tested whether CCN4 stimulates melanoma invasion and metastasis. Wound healing reflects multiple biological processes including cell-matrix and cell-cell interactions, migration, and invasion. Thus wound healing assays were performed to test *Ccn4* knockout on tissue repair in melanoma (Fig.2B). B16F0 cells filled 77.2% of the scratch gap in 24 hours, but the wound healing rate for knockout cells was reduced to 47.0% (F0-KO) and 47.3% (F0-KO’). For B16F10 cells, the wound healing rate was reduced from 76.9% (F10) to 42.9% (F10-KO) and 54.9% (F10-KO’). In migration and invasion assays (Fig.2C and 2D), the migration rate of B16F10 knockout cells was only 51.4% (F10-KO) and 49.3% (F10-KO’) as compared to the parental and control cells, and the invasion rate was also reduced to 41.2% (F10-KO) and 54.9% (F10-KO’), relative to the parental cells.

To assess the invasion and metastasis potential *in vivo*, we performed a spontaneous metastasis assay with subcutaneous injection of B16F0 and its knockout cells (F0-KO) using C57BL/6Ncrl and NOD-scid IL2Rgamma^null^ (NSG) mice. At the experimental endpoint, all mice within a cohort were euthanized and the spontaneous lung metastasis was assayed through *ex vivo* bioluminescence and real time genomic qPCR (17)(11). No metastatic lung colonies or metastatic lung signals were detected using either bioluminescence or qPCR in any of the C57BL/6Ncrl mice. However, spontaneous lung micrometastases from B16F0 in NSG mice were readily revealed by both bioluminescence imaging and qPCR (Fig.2E and 2F). For B16F0-KO cells in NSG mice, the bioluminescent signals for lung micrometastases were lost (Fig.2E) and their metastatic tumor burdens, revealed by genomic qPCR, was repressed by over 80% (Fig.2F). In general, the *in vitro* and *in vivo* results suggested that CCN4 promotes melanoma cell migration, invasion and metastasis, similarly as reported using double nickase-derived variants (11).

### Paracrine CCN4 Stimulates Mouse and Human Melanoma Cell Migration and Invasion

In transwell migration and invasion assays, medium conditioned by mouse fibroblast NIH3T3 was required as chemoattractant for B16 cells (11). Since NIH3T3 releases moderate amounts of CCN4, we created three new NIH3T3 lines with different CCN4 secretion levels to evaluate the importance of paracrine CCN4 signals for melanoma cell migration and invasion (Fig.3A).

**FIGURE 3.**
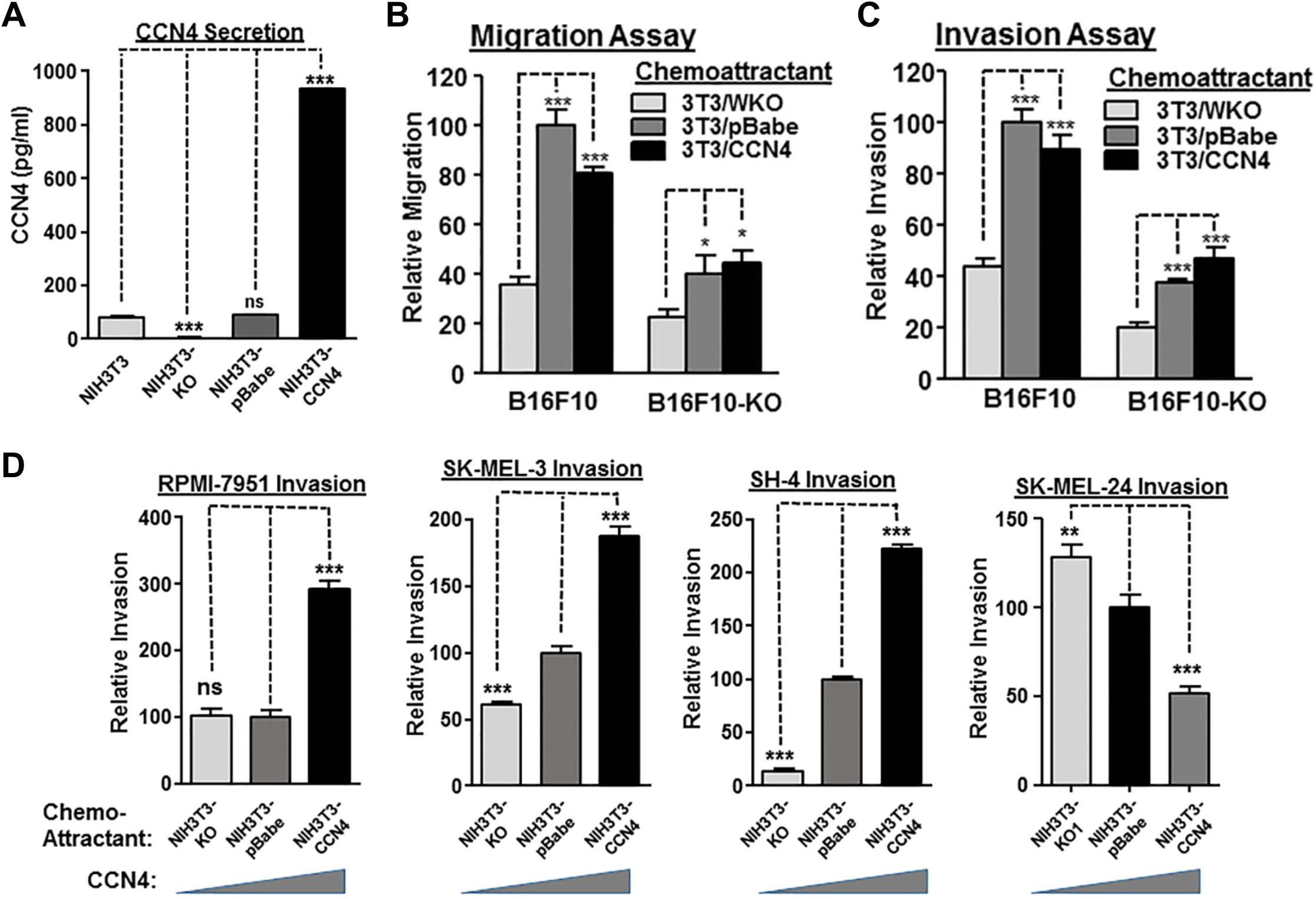
Paracrine CCN4 promotes mouse and human melanoma cell migration and invasion. **A.** CCN4 secretion from knock-out, control, and over-expression cells derived from mouse fibroblast NIH3T3 cells. Culture media were tested by ELISA after cells were plated in serum-free medium for 48 hours. **B.** Migration comparison of B16F10 and B16F10-KO cells. Transwell migration assays were performed in triplicates and serum-free conditioned media from indicated cells was used. **C.** Invasion comparison of B16F10 and B16F10-KO cells. Transwell invasion assays were performed in triplicates and serum-free conditioned media from indicated cells was used. **D.** Cell invasion assays with indicated chemoattractant (conditioned media) using human metastatic melanoma cell lines (RPMI-7951, SK-MEL-3, SH-4 and SK-MEL-24). *: p-value < 0.05; **: p-value < 0.01; ***: p-value < 0.001; and ns: not significant.

In the two assays, the paracrine CCN4 exhibited strong stimulation for both B16F10 and B16F10-KO, especially when CCN4 was present in the conditioned medium (NIH3T3-pBabe) and autocrine CCN4 already sensitized the wild-type B16F10 cells (Fig.3B and 3C). In both cases, the migration and invasion rates were more than doubled, but overexpressed paracrine CCN4 did not further promote migration and invasion (Fig.3B and 3C). For B16F10-KO cells, paracrine CCN4 also promoted migration and invasion, but not as dramatic as exhibited for B16F10 cells, suggesting autocrine CCN4 may help to prime cells for the stimulatory effect from paracrine CCN4. In three out of four human metastatic melanoma cell lines tested, paracrine CCN4 promoted tumor cell invasion as well (Fig.3D). Interestingly, the sensitizing concentration for CCN4 was different for three cells, suggesting a context-dependent cell response to paracrine CCN4 (Fig.3D).

### CCN4 Promotes EMT-like Gene Expression and Represses E-cadherin in B16 Cells

Our previous work showed that CCN4 stimulated invasion and metastasis by promoting EMT-like process within melanoma cells (11). We tested the scenario in our new B16F0 and B16F10 knockout cell pools. In B16F10 cells, *Ccn4* knockout obviously repressed *Snai1*, N-cadherin and vimentin expression at protein level (Fig.4A). While we could not detect E-cadherin by Western Blotting, real-time quantitative RT-PCR revealed that *Ccn4* knockout increased E-cadherin mRNA (*Cdh1*) by more than 50% (75.5% for F10-KO and 50.6% for F10-KO’) (Fig.4B and Supplementary Fig.S1). These results suggest a switch back from mesenchymal to epithelial-like phenotype without CCN4. The EMT marker gene panel we tested with quantitative RT-PCR also showed that *Ccn4* knockout led to decreased expression of *Snai2*, *Zeb2*, N-cadherin (*Cdh2*), vimentin (*Vim*) and fibronectin (*Fn1*) (Fig.4B and Supplementary Fig.S1). The only exception was *Zeb1* mRNA. As observed previously (11), *Zeb1* increased upon *Ccn4* knockout, suggesting it may play a context-specific roles other than promoting EMT in B16 cells. In addition, similar changes at mRNA level for EMT marker gene panel were observed in B16F0 knockout cells (Supplementary Fig.S1).

**FIGURE 4.**
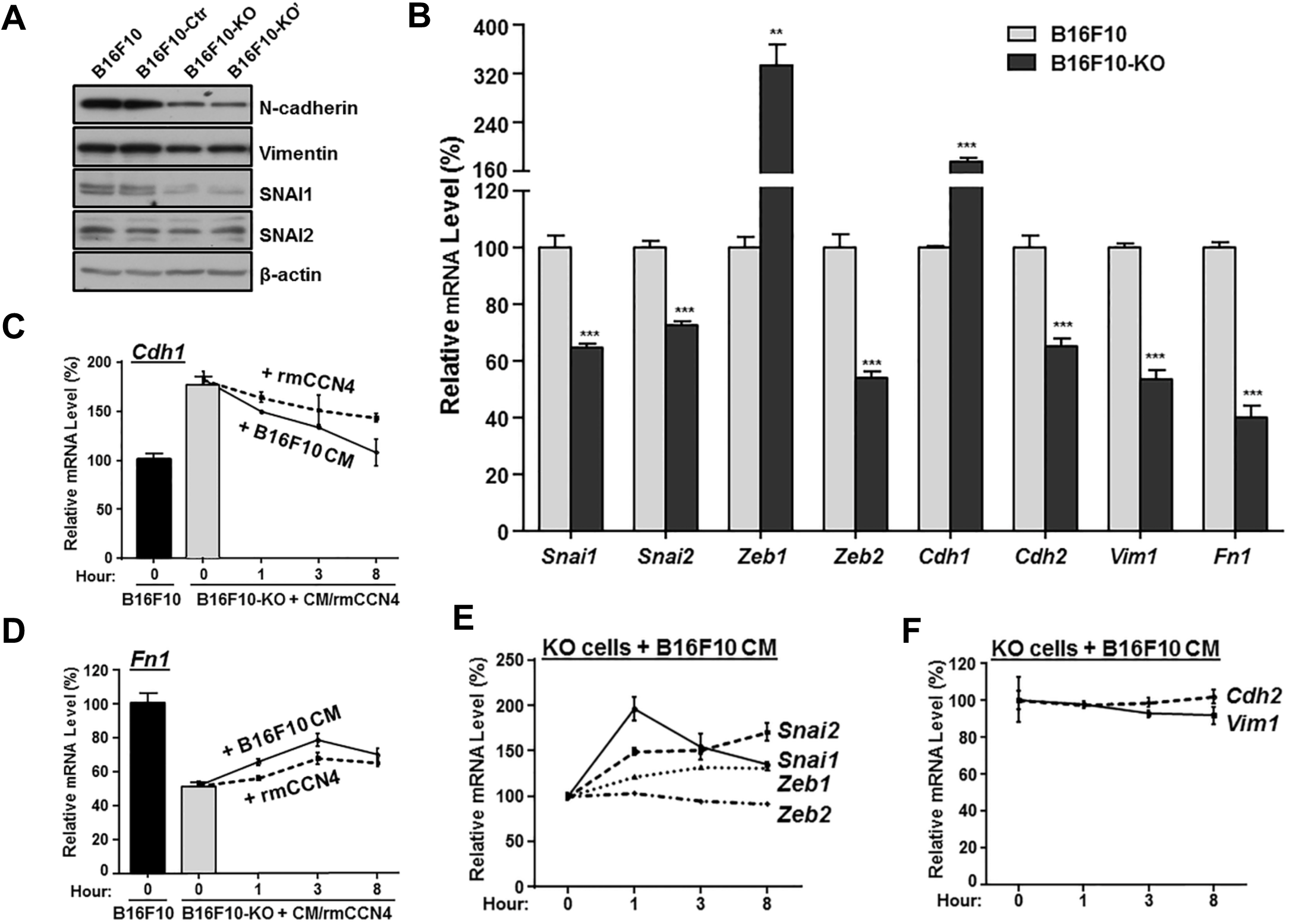
CCN4 promotes EMT gene expression in mouse B16 melanoma cells. **A**. Protein expression comparison of some EMT-associated genes in B16F10 and its knockout cells. Βeta-actin was used as loading control for western blotting. **B**. mRNA expression comparison, as revealed by real-time quantitative RT-PCR, of some EMT-associated transcription factors and EMT marker genes in B16F10 and its knockout cells. **C-F**. mRNA expression change of EMT-associated transcription factors and EMT marker genes in B16F10 knockout cells after CCN4 was supplied in culture medium. Both B16F10 and B16F10-KO cells were plated in complete DMEM with 10% FBS for 48 hours before recombinant mouse CCN4 protein (rmCCN4) was added to B16F10-KO cells (5µg/ml), or B16F10 medium was transferred to B16F10 –KO cells. Cells at indicated time points were harvested for RNA isolation and real-time quantitative RT-PCR. **: p-value < 0.01; ***: p-value < 0.001.

To verify the direct involvement of CCN4 in regulating an EMT-like gene signature, we performed rescue experiments with exogenous CCN4 in the culture medium, using either recombinant mouse CCN4 protein or culture medium from parent B16F10 cells (Fig.4C-4F). Within eight-hours, E-cadherin mRNA (*Cdh1*) was reduced to as low as 78.0% (with rmCCN4, dotted line) and 57.8% (with F10 medium, solid line) of the starting level (Fig.4C). With CCN4 supplementation, we also saw an immediate increase in fibronectin (Fig.4D), *Snai1* and *Snai2* (Fig.4E). Notably, *Snai1*, identified as a major mediator of CCN4-promoted EMT, peaked at the 1 hour time point. In contrast, an increase in *Zeb1* exhibited a time lag in response to CCN4, vimentin trended towards a decrease in time, while N-cadherin and *Zeb2* appeared to be unchanged at any time point (Fig.4E and 4F). The dynamic response in *Snai1* and *Zeb1* suggests a possible negative feedback on *Snai1* expression by *Zeb1*, although the details of this genetic regulatory network remain to be clarified in this context. Nevertheless, these results collectively suggest that CCN4 promotes an EMT-like process in melanoma cells that leads to migration and invasion.

### *Ccn4* Knockout Facilitates B16 Cell Proliferation at Optimal Growth Conditions

Two reports using either mouse K-1735 or human 1205Lu melanoma cells suggest that CCN4 represses melanoma growth in culture medium (8,10). Analogously, we showed previously that *Ccn4* Knockout promoted B16F10 cell growth *in vitro* (11). We then monitored our new B16F0-KO/-KO’ and B16F10-KO/KO’ cells for 2D growth on 96-well plates under optimal growth conditions (Fig.5A). Over three-days, the knockout cells outgrew the parental cells by 23.5% (B16F0-KO) and 20.7% (B16F0-KO’) for B16F0, and 27.7% (B16F10-KO) and 32.8% (B16F10-KO’) for B16F10 cells (Fig.5A). Anchorage-independent growth of knockout cells was also tested using soft agar assays (Fig.5B). With tumor colonies sparsely grown in soft agar for 2 weeks, *Ccn4* knockout increased B16F0 colony formation by 99.7% (F0-KO) and 134% (F0-KO’), while knockout increased B16F10 colony formation by 102% (F10-KO) and 86.0% (F10-KO’). These results showed that disruption of *Ccn4* expression increases melanoma proliferation, and it is consistent with the previous reports (8)(10)(11).

**FIGURE 5.**
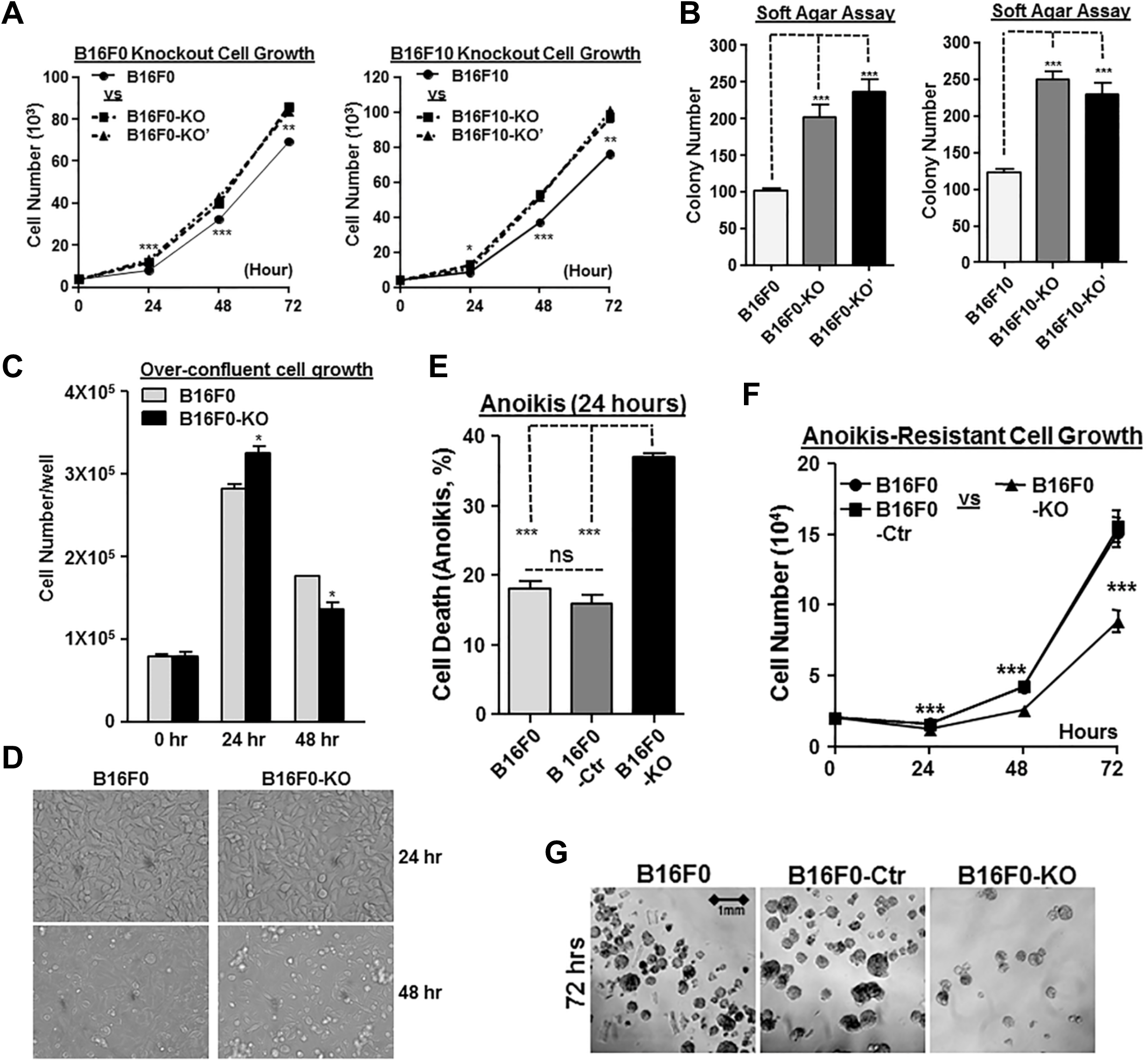
*Ccn4* knockout facilitates B16 cell proliferation at optimal growth conditions. **A.** Growth comparison of B16F0 with its knockout cells (left) and B16F10 with its knockout cells (right) during 3-day period in regular 2d culture. Assays were performed in triplicates on 96-well plates and 2,000 cells were seeded in each well on day 0 and ended before the cells reached full confluence. **B.** Anchorage-independent growth for B16F0 and B16F10 knockout cells. Cells were plated in triplicates in soft agar on 6-well plates. Colonies were fixed and counted after 12 days. **C.** 2-day growth comparison of B16F0 and B16F0-KO cells in overconfluent conditions in 2D culture. Assays were performed in triplicates on 96-well plates and 80,000 cells were seeded in each well on day 0. **D.** Representative photographs from each group in **C**. **E.** Cell death comparison after 24 hours in Anoikis assay. **F.** Cell growth comparison in a total of 72 hours in Anoikis assay. **G.** Representative photographs from each group in **F** at 72 hours. *: p-value < 0.05; ***: p-value < 0.001.

As a transplantable syngeneic mouse model for melanoma, B16 melanoma is known for its rapid growth *in vivo* and the associated problems such as poor vascularization and severe necrosis, which always leads to swift mortality of mice in 3-4 weeks (18). Thus *in vivo*, B16 melanoma are unlikely to experience optimal growth conditions such as oxygen, growth factors and nutrients as provided *in vitro*. In a new 2D growth assay on 96-well plates, we started with 80,000 cells in each well so the cells reached full confluence at the beginning of the assay (Fig.5C). With fresh medium, B16F0-KO still proliferated faster than B16F0 when cell density was assayed after the first 24 hours (Fig.5C), but the medium quality deteriorated at the high cell density and its pH turned more acidic. Between 24 to 48 hours, B16F0-KO cells died at a faster rate than B16F0 (Fig.5D), resulting in a lower number of cells than B16F0 at 48 hours (Fig.5C). The result suggested a much weaker survival response from B16F0-KO against stress after *Ccn4* knockout.

The possible role of CCN4 on the survival of B16 melanoma cells was further investigated using Anoikis assays (Fig.5E-5G). Anoikis is a type of programmed cell death from anchorage-independent cell growth without the support of surrounding extracellular matrix (19). This in vitro assay mimics the growth and survival of the injected tumor cells after the transplantation and during the invasion, circulation, colonialization in metastasis. At the first 24 hours, B16F0-KO cells exhibited a much higher death rate than the B16F0 and B16F0-Ctr cells (Fig.5E, 38.2% vs 19.3% and 17.8%). However, in the next 48 hours, these 3D suspended cells adapted to the new environment and started to exhibit anoikis-resistant, anchorage-independent cell grow (Fig.5F) and eventually formed 3D spheroids (Fig.5G). While we could not compare anoikis-resistant cell growth rate due to the difference on the survival (starting) cell numbers at 24 hours, we did note that CCN4 knockout reduced the cell numbers that survived the Anoikis challenge (24 hour), which resulted in less cells to seed the anoikis-resistant growth stage (48 and 72 hours).

### *Ccn4* Knockout Represses Cell Survival Pathways in B16 Cells

CCN4 activates multiple anti-apoptotic/pro-survival signaling in lung and breast cancer cells, as well as in cardiomyocytes and neuronal cells (20)(21)(22). We then tested the cellular response when B16F0 and -KO cells were treated for 48 hours with 5µM of Mitoxantrone (MTX), a chemotherapy drug that causes DNA damage and induces apoptosis (23). As shown with representative results from each group (Fig.6A), 46.44% of B16F0 cells were still alive (Annexin^−^PI^−^) after treatment, compared with only 21.24% of B16F0-KO cells were living. While both samples contained about 24% of dead cells (Annexin^+^PI^+^), only 24.93% of B16F0 cells were undergoing apoptosis (Annexin^+^PI^−^), compared with 52.99% of dying B16F0-KO cells. Quantitatively, knockout B16F0-KO cells were only half as viable as wild type B16F0 cells (Fig.6B, Annexin^−^PI^−^, 21.6±0.4 and 43.7±3.1, respectively), due to a much stronger apoptotic response upon MTX induction. qPCR analysis revealed that, with *Ccn4* knockout, the expression of two anti-apoptotic factors *Bcl2* and *Mcl1* was reduced in both B16F0 and B16F10 cells (Fig.6C).

**FIGURE 6.**
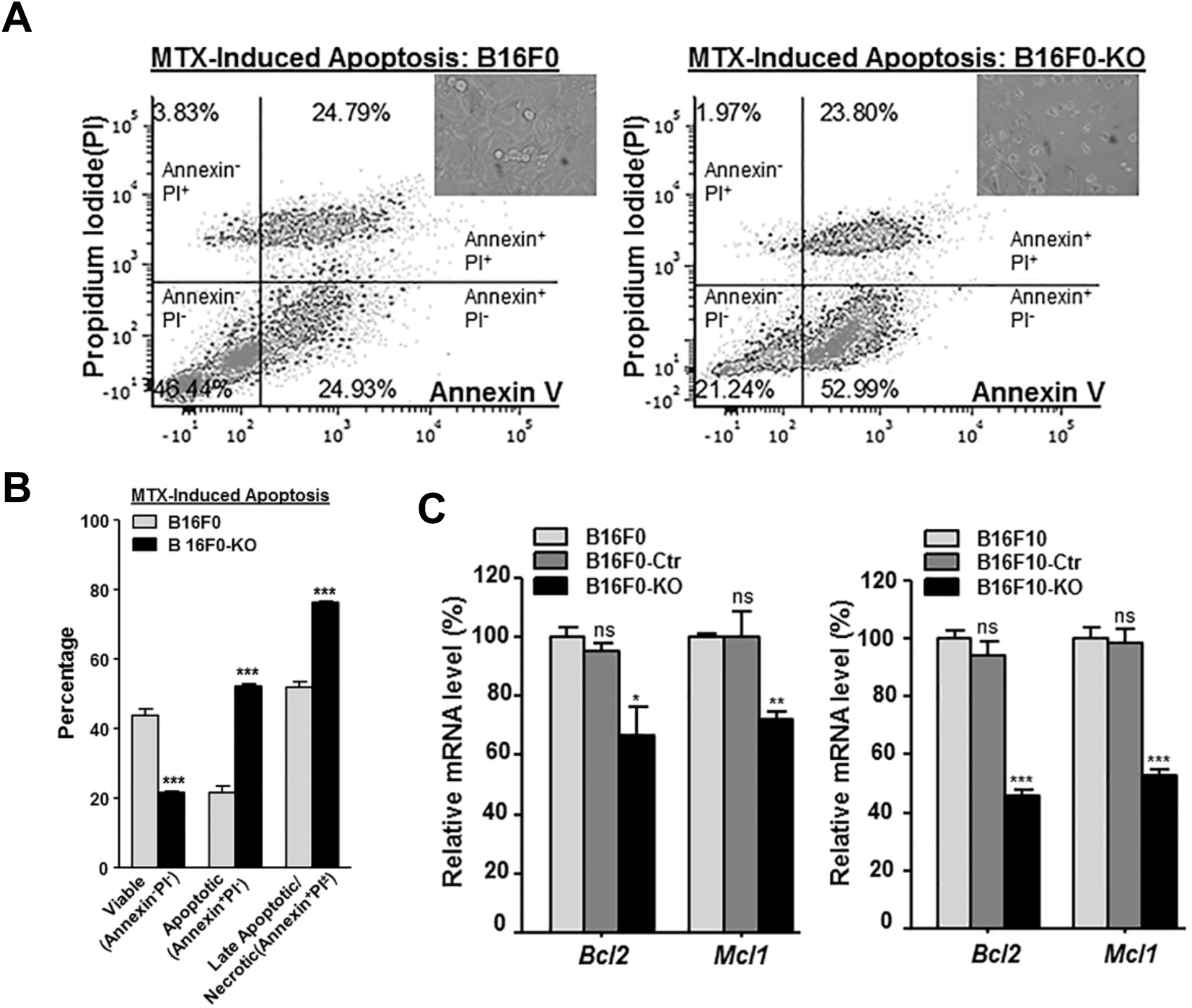
*Ccn4* knockout represses cell survival pathway in B16 cells. **A.** Flow cytometry analysis of mitoxantrone (MTX)-induced apoptosis in B16F0 and B16F0-KO cells. The cells were seeded in triplicates on 6-well plates and treated with MTX for 48 hours before stained for Annexin V and Propidium Iodide (PI). A representative cell distribution from each group are shown. **B.** Comparison of viable portion and apoptotic portion in triplicates of B16F0 and B16F0-KO cells in panel A. **C.** mRNA expression comparison of two anti-apoptotic gene, *Bcl2* and *Mcl1*, in B16F0 and B16F0-KO cells (left), as well as in B16F10 and B16F10-KO cells (right). *: p-value < 0.05; **: p-value < 0.01; ***: p-value < 0.001; and ns: not significant.

### CCN4 shifts B16 cells from a fragile proliferative state to a resilient metastatic state

*In vivo*, mouse K-1735 melanoma clones overexpressing CCN4 grew slower in syngeneic C3H/HeN mice (8), and Shao et al. reported that human 1205Lu melanoma in SCID mice grew faster after the paracrine CCN4 expression was repressed by about 50% in the co-injected stromal fibroblast cells (10). On the other hand, B16F10 *Ccn4* double nickase knockout clones in NSG and C57BL/6Ncrl mice suggested the overall tumor growth was not affected from CCN4 repression *in vivo*, as quantified by comparing the average tumor size for a cohort at a single time point (11). Yet, *in vitro* assays presented in Fig.5 and Fig.6 argued that the net effect of CCN4 on tumor growth could reflect two opposite effects where CCN4 enhances intrinsic survival of tumor cells to environmental stressors and reduces the rate constant associated with cell growth. Environmental stressors include surviving injection into mice and starting tumor growth in a new environment, lack of nutrients due to improper vascularization and angiogenesis (18), and editing by host immunity. To parse these competing effects, we compared the subcutaneous growth dynamics of tumors started by B16F0 variants in immunocompromised (NSG) and immunocompetent (C57BL/6) mice.

First, we observed tumor growth in NSG mice following subcutaneous injection of B16F0 or B16F0-KO cells. Though some tumors from knockout cells seemed smaller initially, the average tumor sizes seemed qualitatively similar between the two groups (Fig.7A, N=5 in each group). Measurable tumors appeared 7 days post injection and were grown until day 21 (Fig.7A). The survival of NSG mice receiving *Ccn4* knockout cells (CCN4 KO) was similar to NSG mice receiving wt B16F0 cells (Fig.7B). Next, we observed tumor growth in immunocompetent C57BL/6Ncrl mice, where CCN4 KO tumors were slower to appear (Fig.7C, N=8 in each group). As wild type B16F0 tumors appeared on day 5 compared to day 7 with CCN4 KO tumors, the wt B16F0 tumors appeared qualitatively bigger than B16F0-KO tumors. A Kaplan–Meier survival estimate showed the *Ccn4* knockout in B16F0 significantly extends survival of C57Bl/6 mice (Fig.7D).

**FIGURE 7.**
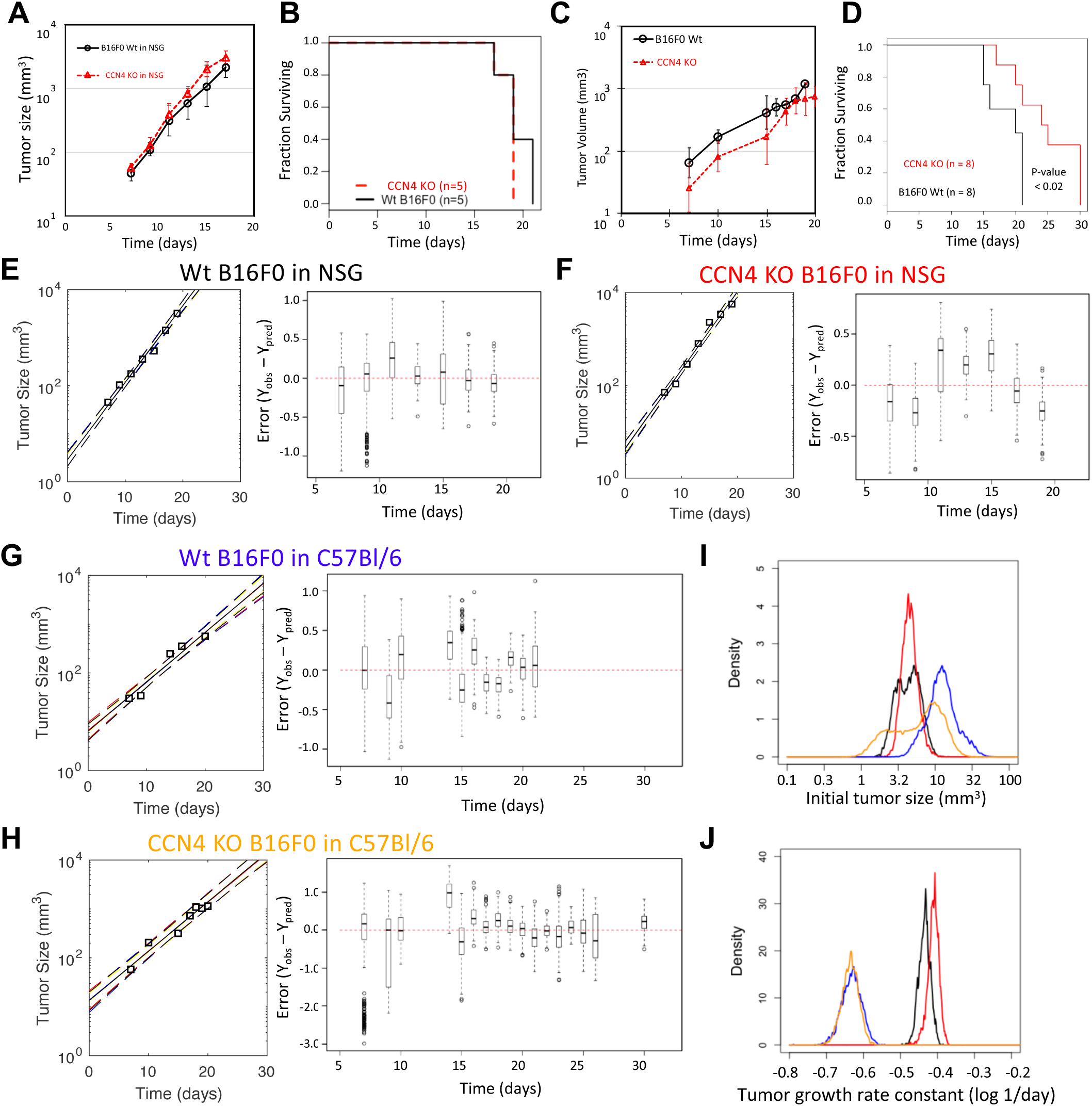
*Ccn4* knockout in B16 cells exhibits different effect on tumor growth in NSG and C57BL/6ncrl mice. Ten NSG mice and sixteen C57BL/6ncrl mice were used in two sets of tumor growth assays (half for B16F0 and half for B16F0-KO cells). Melanoma cells were subcutaneously injected into mice and tumor sizes were recorded from day 7 post-injection until the last day. **A.** Tumor growth from B16F0 and B16F0-KO cells in NSG mice (mean +/-sd). **B.** Kaplan–Meier survival plot of the two groups of NSG mice with B16F0 or B16F0-KO cells (n = 5 in each group). **C.** Tumor growth from B16F0 and B16F0-KO cells in C57BL/6ncrl mice (mean +/-sd). **D.** Kaplan–Meier survival plot of the two groups of C57BL/6ncrl mice with B16F0 or B16F0-KO cells (n = 8 in each group). **E-H.** Modeling the growth of individual tumors for all four groups of mice. Left panels show measurements of tumor growth of one representative mouse (squares) compared to the posterior distribution in the model predictions (solid line corresponds to maximum in the distribution while dotted lines enclose 95% of the distribution). Right panels show box and whisker plots of the distribution in error (Y_obs_ – Y_pred_) of the model fits to all of the animals within a cohort. **I-J.** Comparison of the posterior distributions in the tumor initiation size (T_o_, panel **I**) and growth rate constant (k_p_, panel **J**) for all four groups of mice (B16F0 wt in NSG (black) or in C57BL/6 (gold), CCN4 KO B16F0 in NSG (red) or in C57BL/6 (blue)).

To compare the in vivo growth profiles quantitatively, a mathematical model of exponential tumor growth and an empirical Bayesian approach using an adaptive Markov Chain Monte Carlo (AMCMC) algorithm were used to estimate the initial bolus of tumor cells (T_0_) and the tumor growth rate constant (k_p_) based on the available data (Fig.7E-7H and Supplemental Fig. S2 and S3). Using samples from converged Markov Chains, the posterior distributions in the predicted tumor growth rates were compared to the data for each animal (see Fig.7E-7H and Supplemental Fig.S3). Collectively, the distribution in the error (Y_obs_ – Y_pred_) were centered around zero and were similarly distributed around zero as a function of time, suggesting that the model is of sufficient complexity to capture the observed data. Next, we compared the posterior distributions in the model parameters (Fig.7I and 7J). In NSG mice, the same number of injected wt versus CCN4 KO cells provided a similar initial bolus of tumor cells (median wt 4.41 mm^3^ vs CCN4 KO 4.47 mm^3^, Pearson’s Chi-squared p-value = 1e-05), while k_p_ was greater for CCN4 KO than wt B16F0 cells (median wt 0.368 day^−1^ vs CCN4 KO 0.389 day^−1^, Pearson’s Chi-squared p-value = 1e-05). In contrast, CCN4 KO cells provided a lower initial bolus of tumor cells in C57BL/6 mice compared to wt B16F0 cells despite injecting the same number of cells (median wt 12.2 mm^3^ vs CCN4 KO 6.69 mm^3^, Pearson’s Chi-squared p-value = 1e-05). In addition, k_p_ was slightly lower for CCN4 KO than wt B16F0 cells in C57BL/6 mice (median wt 0.234 day^−1^ vs CCN4 KO 0.232 day^−1^, Pearson’s Chi-squared p-value = 1e-05). In light of these observations, we reanalyzed the in vivo tumor growth measurements for wild type and CCN4 KO variants of B16F10 and YUMM1.7 cells developed using a double nickase CRISPR approach (11). Similarly, we observed that CCN4 KO reduced the initial bolus of tumor cells and increased the tumor growth rate constant, especially in NSG mice (Supplemental Figs. S4-S9). Moreover, the apparent increase in tumor growth rate upon CCN4 KO was consistently reduced in immunocompetent C57BL/6 mice. Collectively, in vivo growth curves suggested that CCN4 shifted cells from a fragile, highly proliferative state to a resilient, low proliferative state. As the results were consistent across cell lines and independent of CRISPR approach, the data suggest that the phenotype is related to CCN4 and not an artifact of the gene editing process. The resilient, low proliferative state also has a propensity for metastasis, as illustrated by Figures 2 and 3. The fragile nature of the CCN4 KO cells was especially apparent in immunocompetent hosts.

To test whether this phenotypic shift induced by CCN4 is also observed in human melanoma, we analyzed RNA-seq data reported by the Cancer Cell Line Encyclopedia (CCLE) for 56 melanoma cell lines. Starting with 56318 genes that had non-zero expression values, we looked for enriched functions associated with those genes that had a significant change in expression correlated with CCN4 expression with an adjusted p-value of less than 0.0005. Genes that appeared upregulated with an increased CCN4 expression (i.e., exhibit a positive correlation coefficient) were queried separately from genes that appeared down-regulated with an increased CCN4 expression (i.e., exhibit a negative correlation coefficient). The correlation and functional enrichment results are summarized in Figure 8 and Supplemental Table S1. For genes that appear to have a positive correlation with CCN4, the UP_KEYWORD: Extracellular Matrix (Benjamini adjusted p-value = 1.3E-21) and the KEGG_PATHWAY: PI3K-Akt Signaling (adj p-value = 1.9E-04) were among the top scoring functional annotations. For genes that appear to have a negative correlation with CCN4, the KEGG_PATHWAY: DNA Replication (adj p-value = 2.30E-06) and the KEGG_PATHWAY: Cell Cycle (adj p-value = 1.30E-2) were among the top scoring functional annotations. Similar to our experimental observations, an increase in CCN4 was associated with reduced expression of DNA replication and cell cycle genes and increased expression of PI3K-Akt signaling and extracellular matrix genes. The increase in CCN4 also corresponded to de-differentiation, as illustrated by a downregulation of MITF and upregulation of ZEB1. Scatter plots illustrate the inherent variability that may dominate findings associated with studying a handful of genes within a small number of cell lines but, as the number of cell lines and genes analyzed increases, the variability averages out to provide more consistent trends.

**FIGURE 8.**
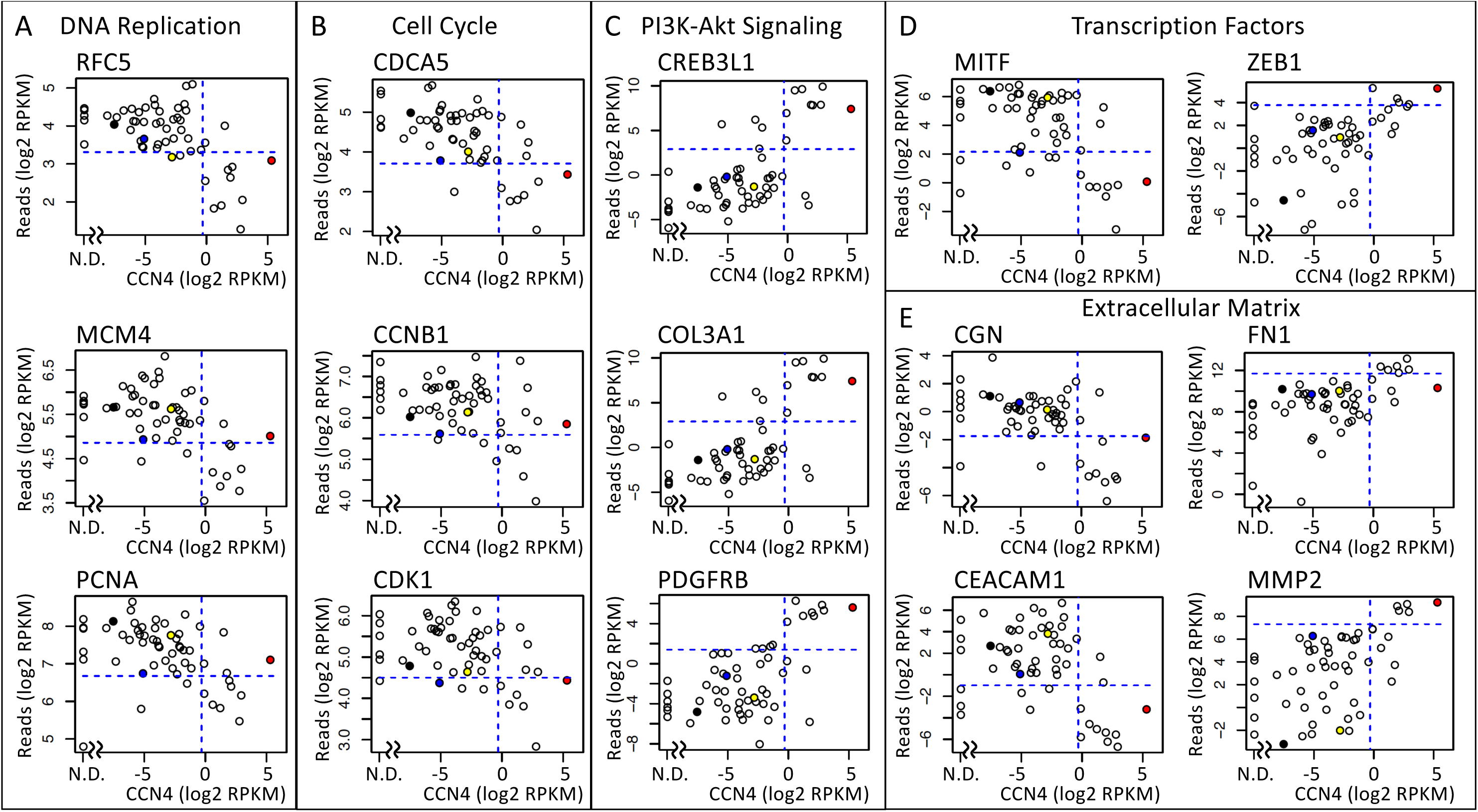
*CCN4* is associated with reduced expression of DNA replication and cell cycle genes and increased expression of PI3K-Akt signaling and extracellular matrix genes in human melanoma cell lines. The expression of a subset of genes associated with DNA replication (**A**), cell cycle (**B**), transcription factors (**C**), PI3K-Akt signaling (**D**), and extracellular matrix (**E**) are plotted against *CCN4* expression assayed by RNA-seq within 56 melanoma cell lines, as reported by the CCLE. Filled circles highlight the values for RPMI-7951 (red), SK-MEL-24 (blue), SK-MEL-3 (black), and SH-4 (yellow) cell lines. Dotted lines indicate the expression threshold used for determining whether the expression of a particular gene is high or low for the Fisher’s exact test. N.D. indicates that *CCN4* was not detected.

## DISCUSSION

For melanoma, revealing pro-metastatic factors within the tumor microenvironment have important implications on tumor diagnosis, prognosis and treatment. Previously, we reported that the secreted matricellular protein CCN4, formerly called WISP1, is a potential invasive melanoma biomarker and therapeutic target (11)(24). Here, we found that CCN4 mRNA expression is associated progressively with reduced overall survival in primary melanoma patients. Using an immunocompetent melanoma model, we generated new *Ccn4*-knockout cell pools using a HDR CRISPR/Cas9 system to generalize results that CCN4 promotes an EMT-like gene signature and stimulates tumor invasion and metastasis (11). Further, we showed that CCN4 enhanced melanoma cell survival and B16 melanoma progression *in vivo*. Though CCN4 repressed B16 tumor cell growth *in vitro* under optimal condition, the effect looked recessive in a set of *in vitro* and *in vivo* assays where tumor cell survival was the dominant factor influencing tumor growth. In short, these data suggest that selecting CCN4 KO clones using a double-nickase CRISPR/Cas9 system did not introduce a phenotypic bias, that is we reject the motivating hypothesis.

CCN4 has been suggested to act as an oncogene to promote tumor cell proliferation, survival, migration, invasion and metastasis in a variety of cancers (13)(14) and as a tumor suppressor in the context of melanoma (8)(10). In light of our data, we note that apparent contradictory observations as to the role of CCN4 in promoting (or, repressing) tumor cell proliferation were more based on observing phenotypic consequences (8)(9)(10)(11)(21)(25)(26)(27)(28). In addition, the experimental design of several studies of CCN4 in the context of melanoma lead to ambiguous interpretations, such as non-physiologic CCN4 overexpression using a strong promoter like CMV, the use of recombinant CCN4 from bacteria, the use of a single cell line or problematic cell lines through the work, and the use of only 2D growth at optimal conditions. Looking back at these data, the possible misinterpretation was highlighted by our results on melanoma cell growth at optimal or sub-optimal stress conditions, *in vitro* or *in vivo* (Fig.5-7). The dominant pro-survival role of CCN4 observed in B16 cells both *in vitro* and *in vivo* was consistent the CCLE data and with the reports in other cellular context in stress conditions (21)(22)(25)(26)(29).

Although the intracellular signaling steps that underpin this shift in phenotype remain unclear, binding of CCN4 with integrins on cell membrane is implicated in the transmembrane signal transduction and intracellular signal activation (30)(31). Further downstream, CCN4 activates numerous signal pathways, including PI3K/AKT and MEK/ERK signaling, that may be responsible for its stimulating effects on cell proliferation, survival, migration and invasion (20)(32)(33). Towards elaborating on the intracellular signaling mechanisms in melanoma cells, we showed that both autocrine and paracrine CCN4 promoted an EMT-like process by altering gene expression in B16 cells. Among the four human melanoma lines tested, only RPMI-7951 secreted a high concentration of CCN4, while the others did not produce detectable CCN4 (11). Nevertheless, three of them migrated towards increased CCN4, suggesting the existence of heterogeneous tumor populations expressing CCN4, or a possible source of CCN4 from tumor stromal cells during invasion. Given the selection advantage of a high proliferation rate in isolating immortal cell lines from human tumor tissue, it is not surprising that human melanoma cell lines that secrete CCN4 are not prevalent. Further work to determine the roles of autocrine and paracrine CCN4 in human melanoma and revealing the exact intracellular molecular pathways in specific cellular context may refine biomolecules that can be therapeutically targeted to limit metastatic progression.

## MATERIALS AND METHODS

### Cell Culture, Conditioned Media and CCN4 ELISA

Mouse melanoma lines B16F0, B16F10, mouse fibroblast line NIH3T3, human metastatic melanoma lines RPMI-7951, SH-4, SK-MEL-3 and SK-MEL-24 were from ATCC and grown as recommended. NIH3T3-derived cells with *Ccn4* knockout (NIH3T3-KO), CCN4 overexpression (NIH3T3-CCN4, formerly NIH3T3-mWisp1) and control retrovirus infection (NIH3T3-pBabe) were described before (11). 48-hour conditioned media (DMEM with 0.1% FBS) from the indicated cells were prepared for transwell assays, and conditioned media with 10% FBS were used for gene expression stimulation. CCN4 concentration in conditioned medium was determined by ELISA using Human WISP-1/CCN4 DuoSet ELISA Development Kit (R&D Systems, Minneapolis, MN).

### Creation of *Ccn4*-Knockout Cell Pools from B16F0 and B16F10 Cells

Mouse *Ccn4* CRISPR/Cas9 KO Plasmid (sc-423705) and Homology-Directed Repair (HDR) plasmids (sc-423705-HDR) were from Santa Cruz Biotechnology (Dallas, Texas). B16F0 cells were transfected with a mix of CRISPR/Cas9 KO plasmid and HDR plasmids, followed by puromycin selection (1.0µg/ml) for 4 days to create the first *Ccn4* knockout cells, B16F0-KO. The cells were expanded, frozen down and passage 3-6 cells were used in this work. To remove the LoxP-flanked puromycin-resistant cassette, B16F0-KO were transfected by a mix of Cre recombinase expression (sc-418923, Santa Cruz Biotechnology) and GFP plasmids. The second *Ccn4* knockout cells, B16F0-KO’, were created by flow sorting of GFP-positive cells. Only passage 4 cells were used in this work. A control cell, B16F0-Ctr, was also made using pBabe-puro retrovirus. The same strategy was used to create B16F10 knockout cells.

### Protein and RNA Analysis

Western blotting was performed as described (11). Mouse monoclonal anti-β-actin (C4) was from Santa Cruz Biotechnology, and the following rabbit monoclonal antibodies were purchased from Cell Signaling Technology (Danvers, MA): anti-Snail (C15D3), anti-Slug (C19G7), anti-Vimentin (D21H3), anti-N-Cadherin (D4R1H).

RNA isolation and qRT-PCR was performed as described (11). Samples for RNA analysis were prepared in biological triplicates and cells were plated on 6-well plates for 48 hours before harvested for gene expression analysis. To induce knockout cells for EMT gene expression, B16F10-KO cells were grown for 24 hours before the medium was replaced by the conditioned medium. The cells were treated for the indicated hours and harvested for RNA isolation. In groups treated with recombinant mouse CCN4 (rmCCN4, 1680-WS-050, R&D Systems), rmCCN4 was added at a final concentration of 5.0µg/ml.

### Functional and Cell Growth Assays

All wound healing, transwell migration, and invasion assays were performed as described (11). 2D cell growth was tested on 96-well plates in triplicate with 2.0×10^3^ cells (optimal condition) or 8.0×10^4^ cells (over-confluent condition) seeded per well as indicated, final cell numbers were measured using ATPlite Luminescence Assay System (Perkin Elmer Inc., Bridgeville, PA). Anchorage-independent cell growth (Soft Agar Assay) was performed on 6-well plates in biological triplicates as described (34). Briefly, 1.0×10^3^ cells were seeded in each well, grown for 2 weeks before the plates were fixed and colonies more than 50 µm in diameter were counted under phase contrast microscopy.

### Anoikis Assay and Chemical Induced Apoptosis

The induced apoptosis in cells without extracellular matrix (ECM) was performed in triplicates as described previously (34). Briefly, 2×10^4^ cells in complete DMEM with 1% methylcellulose (Sigma-Aldrich) were resuspended in six-well low-attachment plates. At indicated time, cells were collected, washed, treated by Accumax (Innovative Cell Technologies) to generate single cell suspensions, and counted after mixed with Trypan Blue. In chemical-induced apoptosis assay, cells in triplicate on 6-well plates were treated with mitoxantrone hydrochloride (MXT, Selleck Chemicals, Houston, TX) for 48 hours at final 5uM. Samples were then stained with FITC Annexin V Apoptosis Detection Kit (BD Biosciences, San Jose, CA), and analyzed by a LSRFortessa flow cytometer (BD Biosciences).

### *In vivo* Tumor Growth/Metastasis Assays, Bioluminescence Imaging and Genomic qPCR

All animal experiments were approved by West Virginia University (WVU) Institutional Animal Care and Use Committee and performed on-site. C57BL/6Ncrl mice (6-8 week-old female) and NOD-scid IL2Rgamma^null^ (NSG, 6-8 week-old male) were from Charles River Laboratories and The Jackson Laboratory, respectively. Mice were injected subcutaneously with 3×10^5^ of indicated cells and tumor sizes were recorded via caliper. Spontaneous lung metastasis were also assayed in tumor-exposed mice following euthanasia using *ex vivo* bioluminescence and genomic qRT-PCR as described (17).

### Modeling and Statistical Analysis

The starting tumor cell population and the intrinsic rate constant for tumor cell growth were determined using the in vivo tumor growth profiles and analyzed quantitatively using an empirical Bayesian approach to estimate the uncertainty associated with the model predictions and parameters, as described previously (35). Briefly, the rate of tumor growth was assumed proportional to size of the tumor (T) in mm^3^ multiplied by a proliferation rate constant (k_p_) and started by injecting tumor cells (i.e., dT/dt = k_p_ * T, where T(t=0) = T_0_). Tumor growth profiles for each animal were analyzed separately with k_p_ assumed to be the same among animals receiving the same cell line variant and with T_0_ unique for each animal. Uncertainty in model predictions and parameters were estimated using an adaptive Markov Chain Monte Carlo (AMCMC) approach. A Pearson’s Chi-squared test for equivalence of distributions was used to assess statistical difference between the posterior distributions in parameter values.

Gene expression and clinical profiles for patients diagnosed with stage I to III melanoma (SKCM) from TCGA were downloaded using the “TCGAbiolinks” (V2.8.2) package in R (V3.5.1). CCLE data was downloaded from the Broad Institute (https://portals.broadinstitute.org/ccle/data Release:081117). Functional enrichment was performed using DAVID v6.8 (https://david.ncifcrf.gov). Kaplan-Meier analysis, Cox proportional hazards modeling, and Fisher’s exact tests were performed using the “survival” (V2.42-6), “survminer” (V0.4.2), and “stats” (V3.5.1) packages in R. Unpaired Student’s t-test (two-tailed) or one-way ANOVA followed by Tukey’s multiple comparison ad hoc post-test were performed with GraphPad Prism (version 5). Results shown as mean±s.d. A p-value of <0.05 was considered statistically significant.

## ACKNOWLEDGEMENTS

This work was supported by National Science Foundation (NSF CBET-1644932 to DJK) and National Cancer Institute (NCI 1R01CA193473 to DJK). The content is solely the responsibility of the authors and does not necessarily represent the official views of the NSF or NCI. Small animal imaging and image analysis were performed in the West Virginia University Animal Models & Imaging Facility, which has been supported by the West Virginia University Cancer Institute and NIH grants P20 RR016440 and P30RR032138/P30GM103488.

## AUTHOR CONTRIBUTIONS

Conceptualized study: WD and DJK; performed experiments: WD, AF, and SLM; analyzed data: WD, AF, and DJK; and drafted original manuscript: WD and DJK. All authors edited and approved of the final version of the manuscript.

The authors declare that they have no conflicts of interest with the contents of this article.

