## Supplemental Table S1 for "Cell Communication Network factor 4 (CCN4/WISP1) shifts melanoma cells from a fragile proliferative state to a resilient metastatic state"

Supplemental Table S1. Functional annotation of genes significantly correlated with CCN4 expression in 56 melanoma cell lines associated with the Cancer Cell Line Encyclopedia. Adjusted p-values are reported for functional annotation. A Fisher Exact test p-value adjusted for multiple hypothesis testing indicated whether gene expression was differentially associated with an increase in CCN4 expression. DAVID (http://david.ncifcrf.gov) was used for functional annotation.

KEGG_PATHWAY: DNA Replication
BH Adj p-value 2.30E-06

| Gene_Symbol | Adj Fisher Exact P-value | Correlation w/ CCN4 |
|---|---|---|
| FEN1 | 1.11E-04 | -0.239 |
| MCM3 | 1.42E-06 | -0.239 |
| MCM4 | 1.61E-05 | -0.233 |
| MCM6 | 3.46E-04 | -0.188 |
| MCM7 | 1.42E-06 | -0.285 |
| PCNA | 1.36E-04 | -0.195 |
| POLA1 | 1.42E-06 | -0.176 |
| POLD1 | 1.11E-04 | -0.145 |
| POLD2 | 3.46E-04 | 0.017 |
| POLE | 2.43E-04 | -0.261 |
| PRIM1 | 3.69E-07 | -0.257 |
| RFC2 | 4.42E-06 | -0.291 |
| RFC3 | 4.42E-06 | -0.259 |
| RFC4 | 4.54E-05 | -0.201 |
| RFC5 | 3.69E-07 | -0.289 |
| RPA2 | 1.61E-05 | -0.182 |
| RPA3 | 3.46E-04 | -0.102 |
| SSBP1 | 3.33E-04 | -0.205 |

KEGG_PATHWAY: Cell Cycle
BH Adj p-value 1.30E-02

| Gene_Symbol | Adj Fisher Exact P-value | Correlation w/ CCN4 |
|---|---|---|
| CCNB1 | 2.23E-02 | -0.181 |
| CCNB2 | 1.00E-02 | -0.112 |
| CCNE1 | 4.41E-05 | -0.233 |
| CDC25A | 4.66E-06 | -0.272 |
| CDC45 | 3.69E-07 | -0.256 |
| CDC6 | 1.42E-06 | -0.291 |
| CDC7 | 3.46E-04 | -0.224 |
| CDCA5 | 1.06E-05 | -0.290 |
| CDCA7 | 1.61E-05 | -0.142 |
| CDK1 | 2.07E-03 | -0.247 |
| CDK2 | 4.41E-05 | -0.189 |
| CDK4 | 3.33E-04 | -0.109 |
| CDK5 | 4.54E-05 | -0.199 |
| CUL1 | 3.33E-04 | -0.212 |
| DBF4 | 4.42E-06 | -0.283 |
| E2F1 | 1.11E-04 | -0.219 |
| E2F2 | 4.54E-05 | -0.231 |
| E2F3 | 3.46E-04 | -0.226 |
| EP300 | 4.66E-06 | -0.168 |
| ESPL1 | 1.42E-06 | -0.339 |
| HDAC1 | 1.36E-04 | -0.183 |
| MAD1L1 | 4.54E-05 | -0.116 |
| MCM3 | 1.42E-06 | -0.239 |
| MCM4 | 1.61E-05 | -0.233 |
| MCM5 | 2.07E-03 | -0.185 |
| MCM6 | 3.46E-04 | -0.188 |
| MCM7 | 1.42E-06 | -0.285 |
| ORC1 | 4.42E-06 | -0.242 |
| ORC5 | 1.36E-04 | -0.250 |
| ORC6 | 1.11E-04 | -0.311 |
| PCNA | 1.36E-04 | -0.195 |
| PKMYT1 | 4.41E-05 | -0.281 |
| PTTG1 | 1.96E-08 | -0.253 |

Transcription Factors and Cell interaction Genes

| Gene_Symbol | Adj Fisher Exact P-value | Correlation w/ CCN4 |
|---|---|---|
| CEACAM1 | 2.84E-05 | -0.147 |
| CGN | 3.69E-07 | -0.130 |
| MITF | 4.89E-04 | -0.255 |
| NRCAM | 7.67E-04 | -0.175 |
| CDH11 | 4.54E-05 | 0.180 |
| FN1 | 1.42E-05 | 0.233 |
| VIM | 1.07E-02 | 0.411 |
| ZEB1 | 1.21E-04 | 0.678 |

KEGG_PATHWAY: PI3K-Akt Signaling
BH Adj p-value 1.90E-04

| Gene_Symbol | Adj Fisher Exact P-value | Correlation w/ CCN4 |
|---|---|---|
| CHRM2 | 1.36E-04 | 0.042 |
| COL1A1 | 1.36E-04 | 0.151 |
| COL1A2 | 2.18E-06 | 0.629 |
| COL3A1 | 3.46E-04 | 0.176 |
| COL5A1 | 3.33E-04 | 0.166 |
| COL6A1 | 2.44E-09 | 0.077 |
| COL6A2 | 1.36E-04 | 0.294 |
| COL6A3 | 1.42E-06 | 0.308 |
| COMP | 4.66E-06 | 0.233 |
| CREB3L1 | 8.31E-08 | 0.404 |
| EGFR | 7.67E-04 | 0.135 |
| FGF2 | 1.61E-05 | 0.169 |
| FGF5 | 2.18E-06 | 0.106 |
| FGF7 | 1.11E-04 | 0.177 |
| FN1 | 1.11E-04 | 0.365 |
| GNGT2 | 1.21E-04 | 0.370 |
| HGF | 4.41E-05 | 0.041 |
| IL6 | 7.67E-04 | 0.025 |
| ITGA11 | 1.36E-04 | 0.463 |
| ITGA5 | 1.21E-04 | 0.305 |
| ITGA8 | 1.61E-05 | 0.133 |
| ITGAV | 9.09E-04 | 0.290 |
| ITGB1 | 3.69E-07 | 0.367 |
| JAK1 | 1.36E-04 | 0.190 |
| LAMB2 | 4.54E-05 | 0.115 |
| LAMC2 | 9.11E-05 | 0.968 |
| LPAR1 | 9.11E-05 | 0.301 |
| MAPK3 | 3.46E-04 | 0.113 |
| NGF | 4.54E-05 | 0.130 |
| PDGFC | 1.61E-05 | 0.372 |
| PDGFRA | 1.19E-05 | 0.283 |
| PDGFRB | 4.54E-05 | 0.538 |
| TEK | 4.41E-05 | 0.148 |
| THBS1 | 1.21E-04 | 0.194 |
| TNXB | 9.11E-05 | 0.114 |
| VEGFC | 1.96E-08 | 0.354 |

UP_KEYWORD: Extracellular Matrix
BH Adj p-value 1.30E-21

| Gene_Symbol | Adj Fisher Exact P-value | Correlation w/ CCN4 |
|---|---|---|
| ADAMTS1 | 1.61E-05 | 0.148 |
| ADAMTS10 | 1.36E-04 | 0.151 |
| ADAMTS12 | 2.18E-06 | 0.629 |
| ADAMTS2 | 3.46E-04 | 0.176 |
| ADAMTS5 | 3.33E-04 | 0.166 |
| ADAMTS6 | 2.44E-09 | 0.077 |
| ADAMTS7 | 1.36E-04 | 0.294 |
| ADAMTSL1 | 1.42E-06 | 0.308 |
| ADAMTSL4 | 4.66E-06 | 0.233 |
| ASPN | 1.16E-07 | 0.215 |
| BGN | 1.61E-05 | 0.262 |
| CCDC80 | 2.84E-05 | 0.191 |
| COL12A1 | 4.66E-06 | 0.209 |
| COL1A1 | 1.61E-05 | 0.280 |
| COL1A2 | 1.11E-04 | 0.365 |
| COL3A1 | 4.42E-06 | 0.270 |
| COL5A1 | 4.66E-06 | 0.395 |
| COL6A1 | 4.66E-06 | 0.656 |
| COL6A2 | 9.87E-07 | 0.691 |
| COL6A3 | 4.54E-05 | 0.172 |
| COL8A1 | 3.33E-04 | 0.111 |
| COMP | 1.42E-06 | 0.171 |
| CPZ | 4.54E-05 | 0.502 |
| CTGF | 1.61E-05 | 0.163 |
| DCN | 4.54E-05 | 0.115 |
| DPT | 9.11E-05 | 0.968 |
| ECM2 | 1.36E-04 | 0.358 |
| EFEMP1 | 1.11E-04 | 0.162 |
| ELN | 1.06E-05 | 0.179 |
| FBLN2 | 4.42E-06 | 0.291 |
| FBN1 | 3.69E-07 | 0.323 |
| FBN2 | 9.09E-04 | 0.117 |
| FN1 | 1.42E-05 | 0.245 |
| FREM3 | 7.67E-04 | 0.040 |
| HSPG2 | 9.11E-05 | 0.114 |
| LAMB2 | 1.61E-05 | 0.382 |
| LAMC2 | 1.42E-06 | -0.023 |
| LOXL2 | 8.98E-04 | 0.278 |
| LTBP2 | 4.42E-06 | 0.320 |
| LUM | 9.11E-05 | 0.227 |
| MFAP4 | 9.87E-07 | 0.573 |
| MFAP5 | 1.06E-05 | 0.210 |
| MMP19 | 1.42E-06 | 0.443 |
| MMP2 | 1.21E-04 | 0.725 |
| NID2 | 8.31E-08 | 0.105 |
| PODN | 1.06E-05 | 0.187 |
| PODNL1 | 3.46E-04 | 0.496 |
| POSTN | 1.11E-04 | 0.199 |
| PXDN | 1.19E-05 | 0.362 |
| SPOCK1 | 6.55E-04 | 0.176 |
| SPON1 | 4.54E-05 | 0.189 |
| TNXB | 3.69E-07 | 0.144 |
| VCAN | 6.55E-04 | 0.336 |
| WNT2 | 1.42E-06 | 0.101 |
| WNT5B | 4.42E-06 | 0.313 |
