## Supplemental Figure S1 for "Cell Communication Network factor 4 (CCN4/WISP1) shifts melanoma cells from a fragile proliferative state to a resilient metastatic state"

A

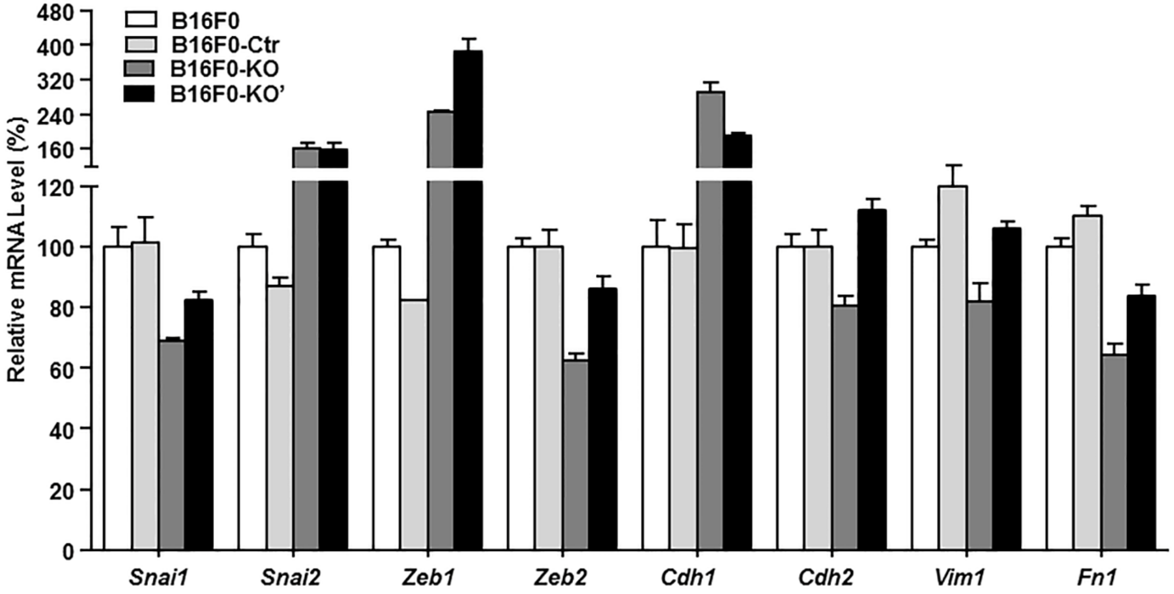


B

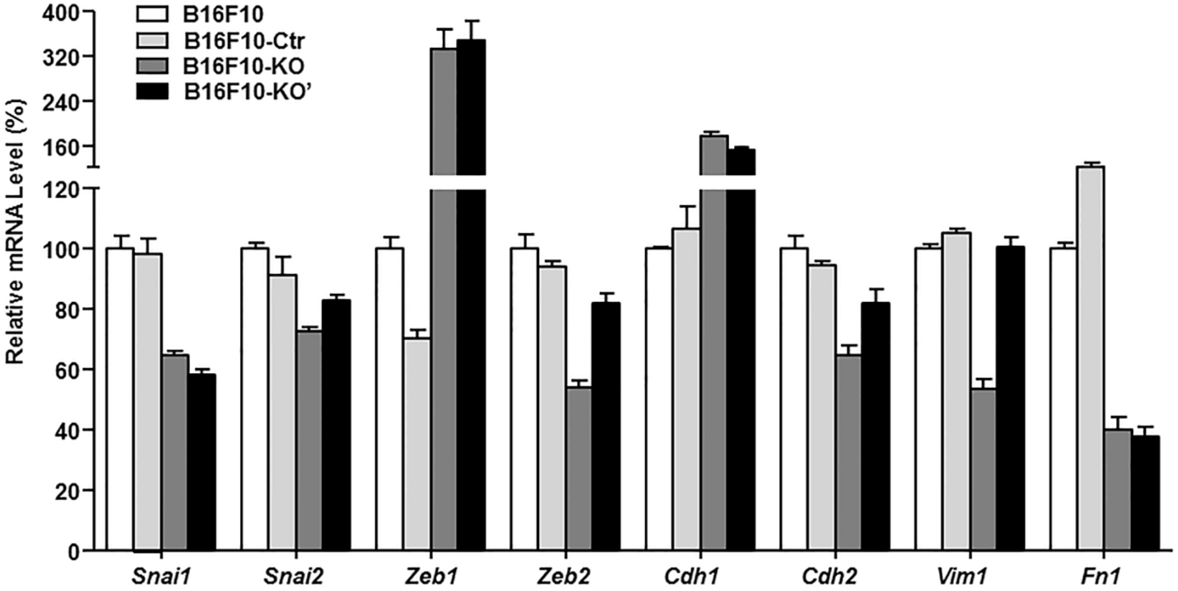


**Supplementary Figure S1. Knockout of *Ccn4* in mouse melanoma B16 cells represses EMT gene expression.** mRNA expression of certain EMT-associated transcription factors and EMT marker genes were compared among two knockout cells (-KO and KO') and control cells (-Ctr) from B16F0 (**A**) and B16F10 (**B**).
