## Supplemental Figure S3 for "Cell Communication Network factor 4 (CCN4/WISP1) shifts melanoma cells from a fragile proliferative state to a resilient metastatic state"

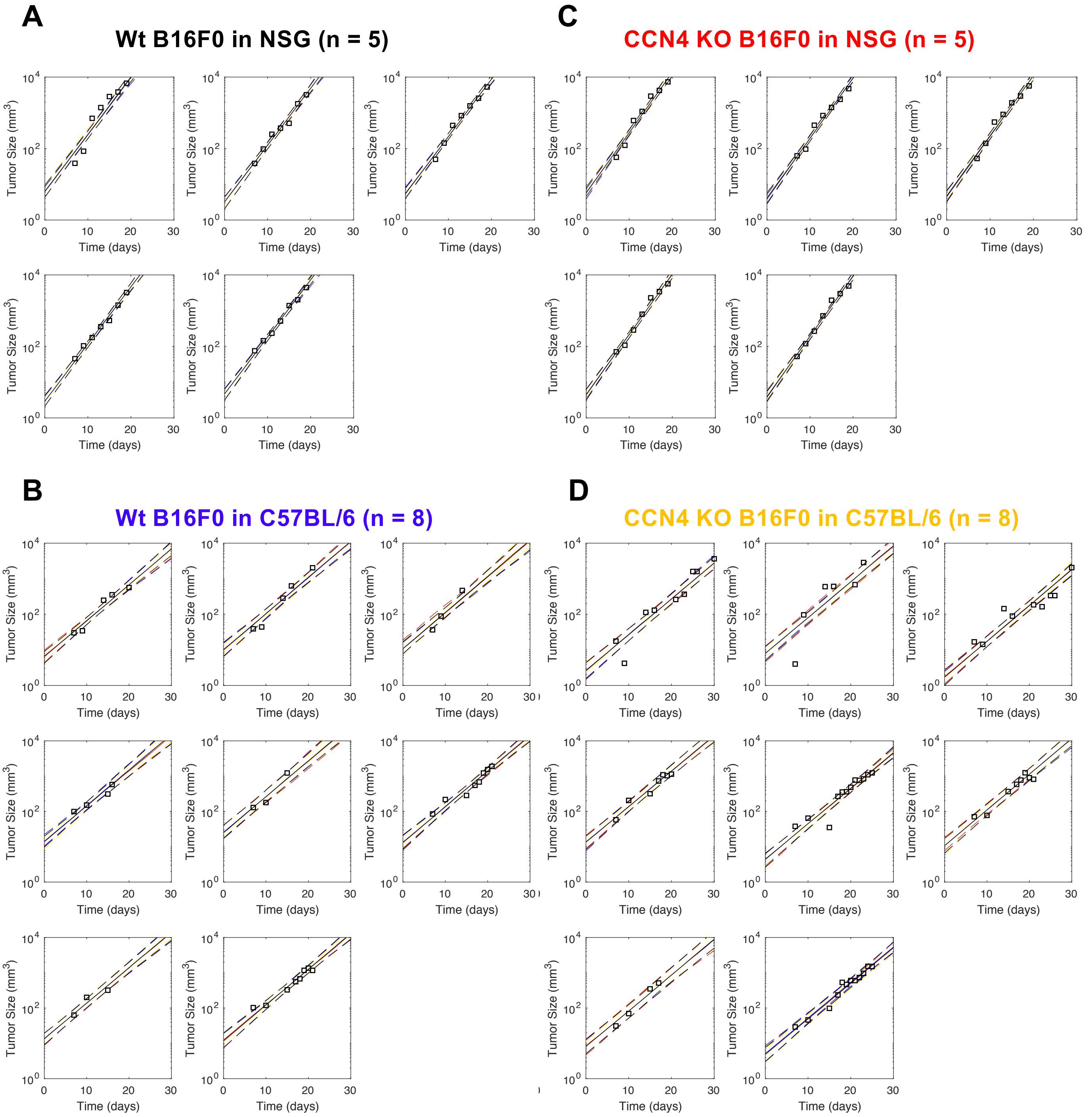


**Supplementary Figure S3. The tumor growth modeling in mice with subcutaneous injection of B16F0 or its knockout cells.** Measures of tumor size at different days (squares) was compared to the posterior distribution in the model predictions (solid line corresponds to maximum in the distribution while dotted lines enclose 95% of the distribution) for each mouse in the cohort. The four groups were B16F0 cells in NSG mice (A, n = 5) or in C57BL/6Ncrl mice (B, n = 8) and *Ccn4* KO B16F0 cells in NSG mice (C, n = 5) or in C57BL/6Ncrl mice (D, n = 8).
