## Supplemental Figure S2 for "Cell Communication Network factor 4 (CCN4/WISP1) shifts melanoma cells from a fragile proliferative state to a resilient metastatic state"

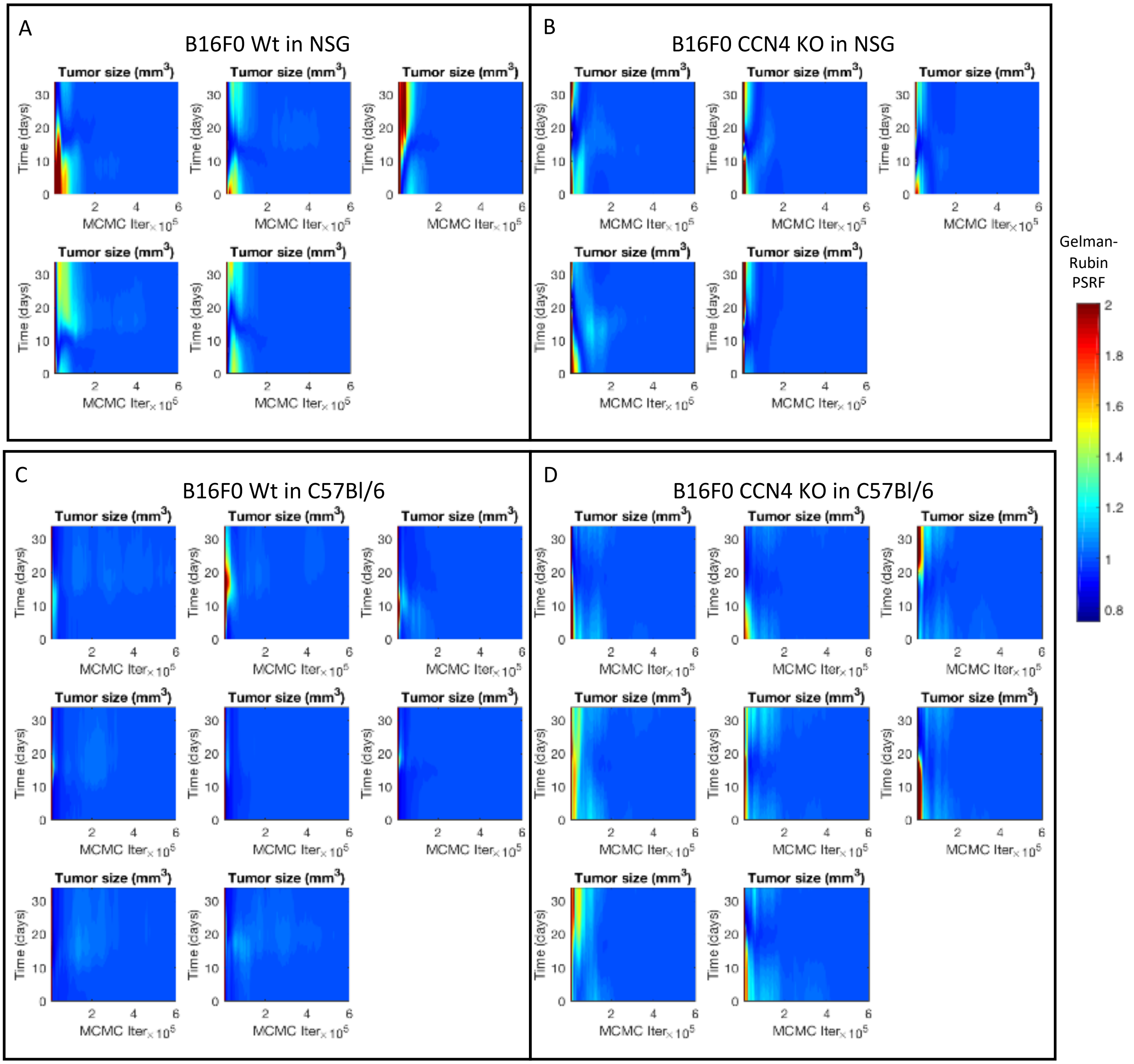


**Supplementary Figure S2. Convergence of AMCMC results for the tumor growth model.** A contour plot of the Gelman-Rubin potential scale reduction factor (PSRF) statistic of the model predictions shown as functions of time (i.e., the y-axis) and AMCMC step (i.e., the x-axis). Three parallel chains were used to calculate the Gelman-Rubin statistics for the model-based inference of the observed tumor growth following subcutaneous implantation of wt B16F0 cells (panels A and C) or B16F0 *Ccn4* (*Wisp1*) KO cells (panels B and D) in either NSG (panels A and B) or C57BL/6 (panels C and D) mice. Each panel corresponds to the convergence to the observed tumor growth in each mouse within the cohort. Values of the PSRF less than 1.2 suggest convergence of the chains.
