## Supplemental Figure S4 for "Cell Communication Network factor 4 (CCN4/WISP1) shifts melanoma cells from a fragile proliferative state to a resilient metastatic state"

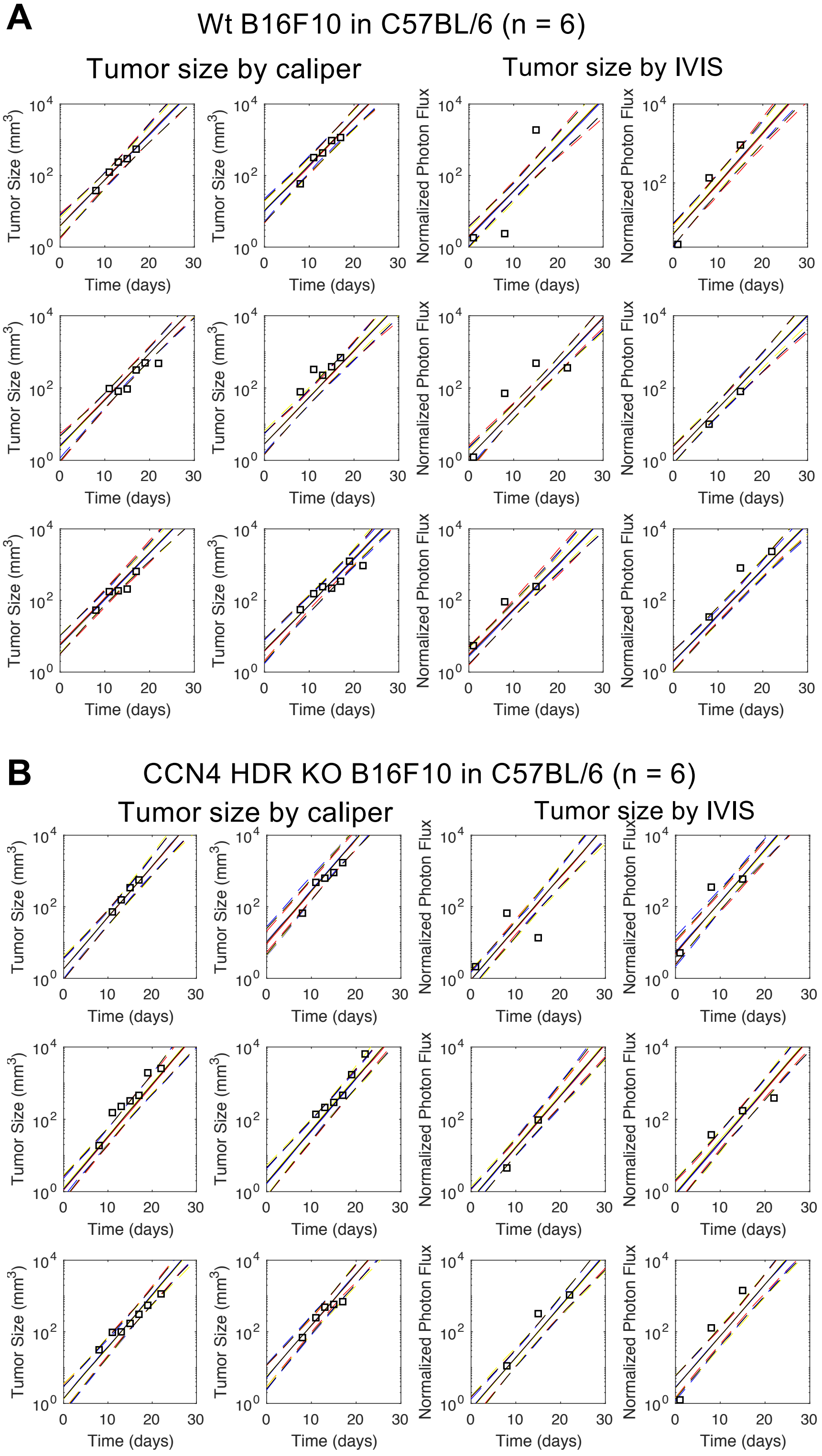


**Supplementary Figure S4. The tumor growth modeling in mice with subcutaneous injection of B16F10 or its HDR knockout cells.** Measures of tumor size obtained using calipers (left 2 columns) or IVIS (right two columns) at different days (squares) was compared to the posterior distribution in the model predictions (solid line corresponds to maximum in the distribution while dotted lines enclose 95% of the distribution) for each mouse in the cohort. The two groups were wild type B16F10 cells (A, n = 6) and *Ccn4* HDR KO B16F10 cells (B, n = 6) in C57BL/6Ncrl mice.
