## Supplemental Figure S5 for "Cell Communication Network factor 4 (CCN4/WISP1) shifts melanoma cells from a fragile proliferative state to a resilient metastatic state"

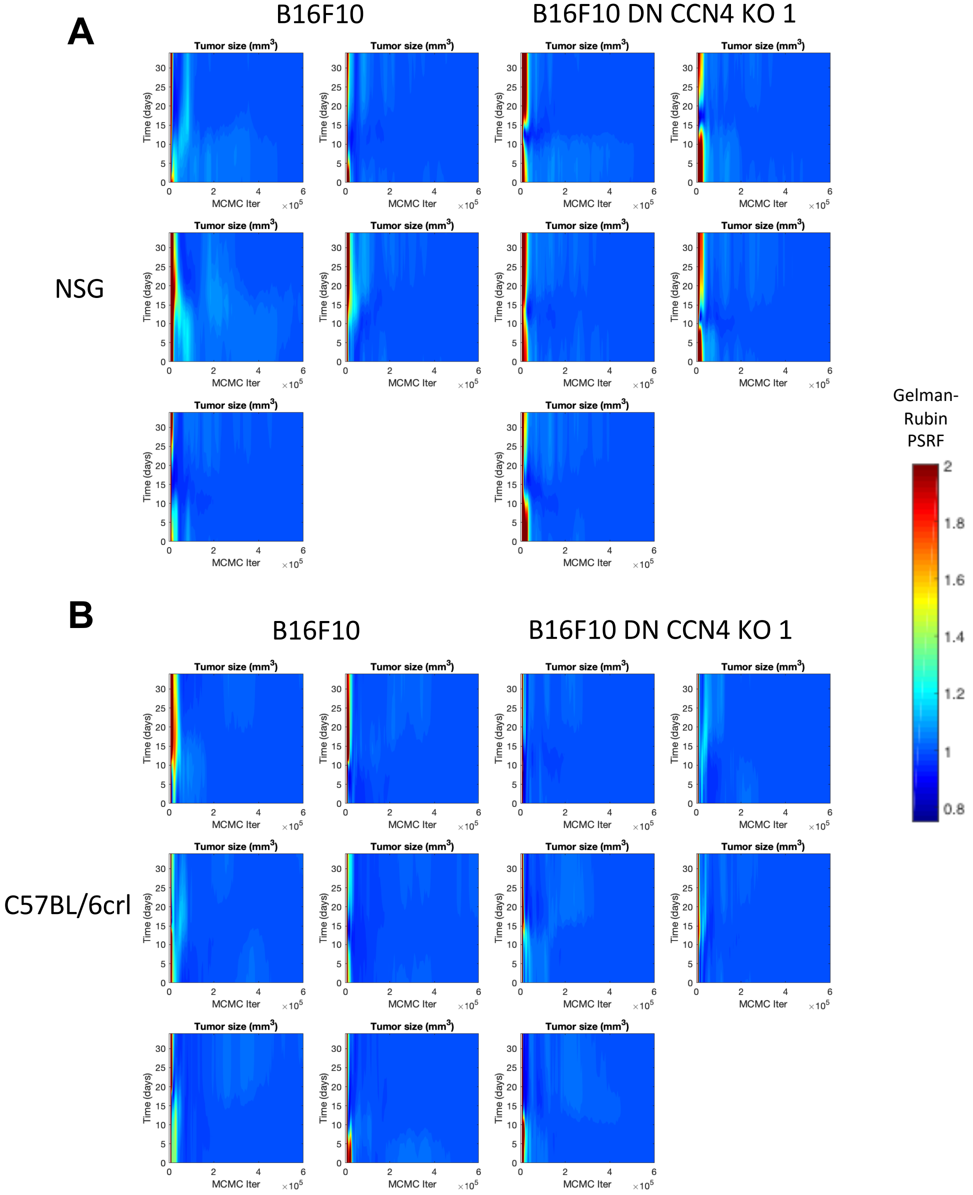


**Supplementary Figure S5. Convergence of AMCMC results for the B16F10 WT/DN KO tumor growth model.** The AMCMC algorithm used an unbiased proposal distribution that was scaled by a covariance matrix estimated from a learning period of 50,000 steps. The proposed steps within parameter space were evaluated using a Metropolis-Hastings algorithm with a targeted acceptance fraction equal to 0.2. Using three independent chains, the Gelman-Rubin Potential Scale Reduction Factor was applied to the model predictions to test for convergence of the chains. A contour plot of the Gelman-Rubin potential scale reduction factor (PSRF) statistic of the model predictions shown as functions of time (i.e., the y-axis) and AMCMC step (i.e., the x-axis). The panels show the Gelman-Rubin statistics for the model-based inference of the observed tumor growth following subcutaneous implantation of wt B16F10 cells (left subpanels) or B16F10 *Ccn4* (*Wisp1*) Double Nickase KO cells (right subpanels) in either NSG (panel A) or C57BL/6crl (panel B) mice. Each panel corresponds to the convergence to the observed tumor growth in each mouse within the cohort. Values of the PSRF less than 1.2 suggest convergence of the chains. The converged segments of the AMCMC chains, which each contained $3.5x10^5$ AMCMC steps, represent samples drawn from the posterior distributions in the model predictions and parameters.
