## Supplemental Figure S6 for "Cell Communication Network factor 4 (CCN4/WISP1) shifts melanoma cells from a fragile proliferative state to a resilient metastatic state"

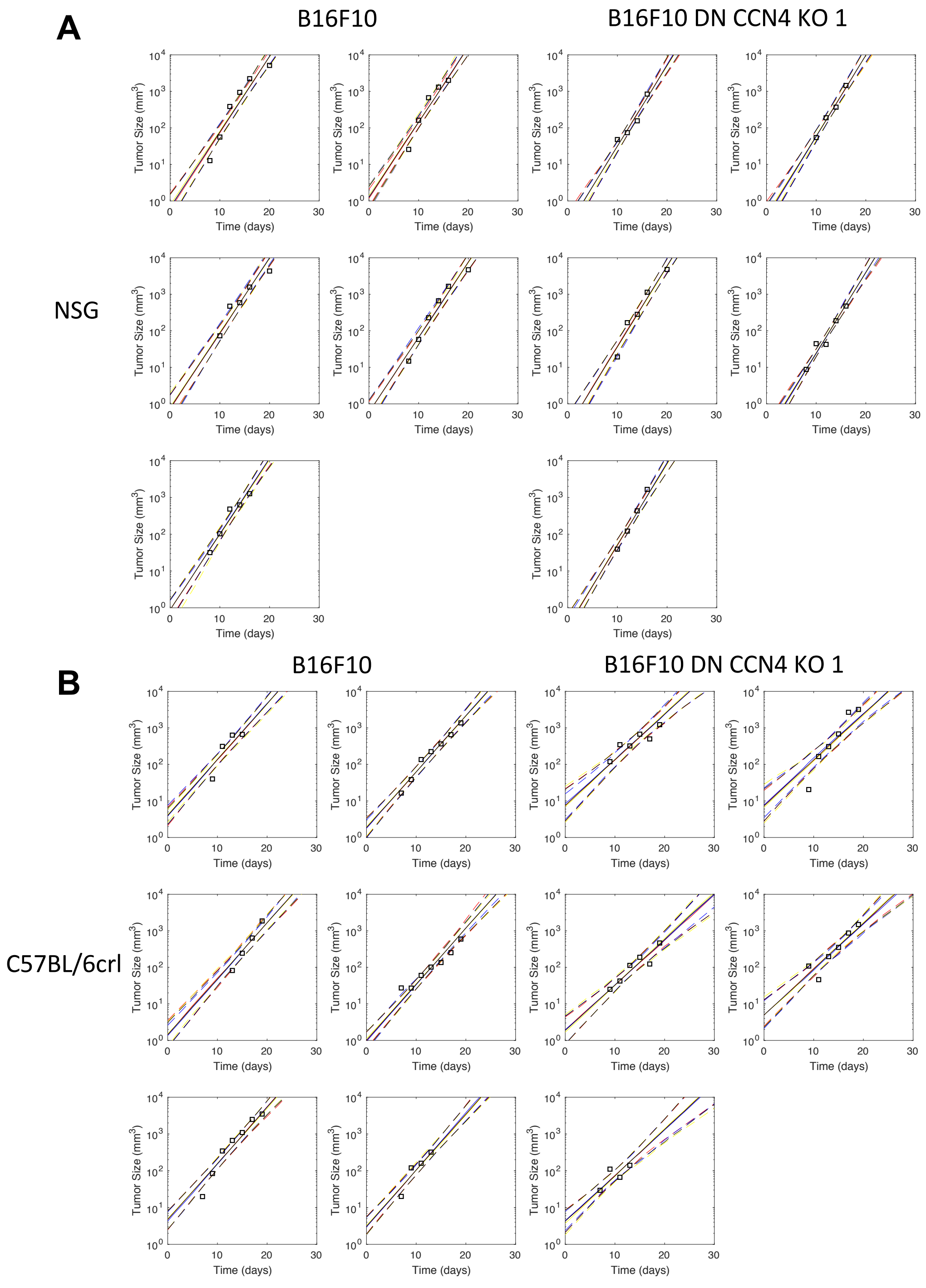


**Supplementary Figure S6. The tumor growth modeling in mice with subcutaneous injection of B16F10 or its *Ccn4* Double Nickase knockout cells.** Measures of tumor size at different days (squares) was compared to the posterior distribution in the model predictions (solid line corresponds to maximum in the distribution while dotted lines enclose 95% of the distribution) for each mouse in the cohort. B16F0 cells (left subpanels) and and *Ccn4* KO B16F0 cells (right subpanels) were injected into NSG mice (A) or into C57BL/6Ncrl mice (B).
