## Supplemental Figure S8 for "Cell Communication Network factor 4 (CCN4/WISP1) shifts melanoma cells from a fragile proliferative state to a resilient metastatic state"

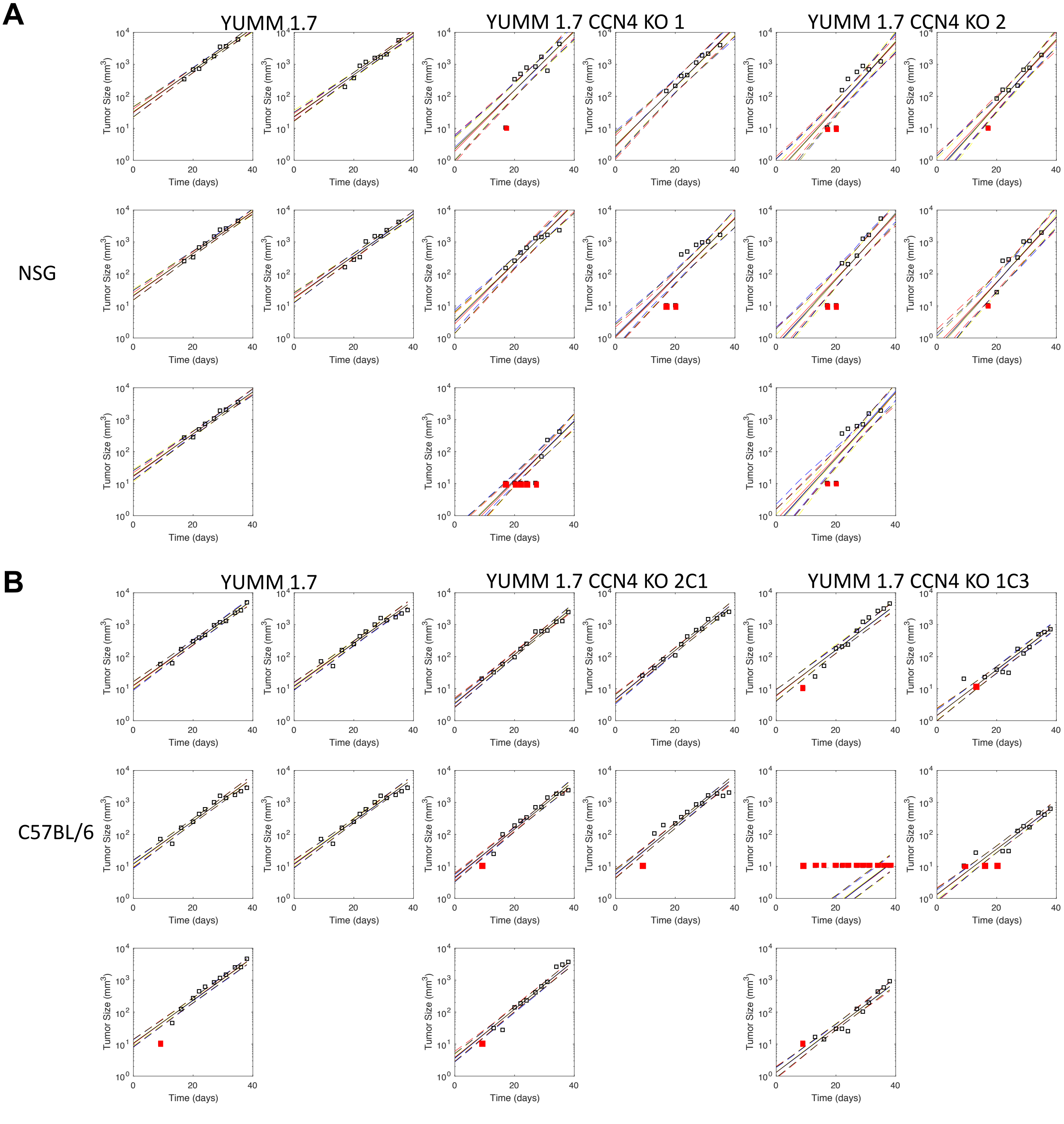


**Supplementary Figure S8. The tumor growth modeling in mice with subcutaneous injection of YUMM1.7 or its *Ccn4* Double Nickase knockout cells.** Measures of tumor size at different days (squares) was compared to the posterior distribution in the model predictions (solid line corresponds to maximum in the distribution while dotted lines enclose 95% of the distribution) for each mouse in the cohort. YUMM1.7 cells (left subpanels) and and *Ccn4* KO YUMM1.7 cells (midle and right subpanels) were injected into NSG mice (A) or into C57BL/6Ncrl mice (B). Red squares indicate that the tumor was too small to be measured.
