## Supplemental Figure S9 for "Cell Communication Network factor 4 (CCN4/WISP1) shifts melanoma cells from a fragile proliferative state to a resilient metastatic state"

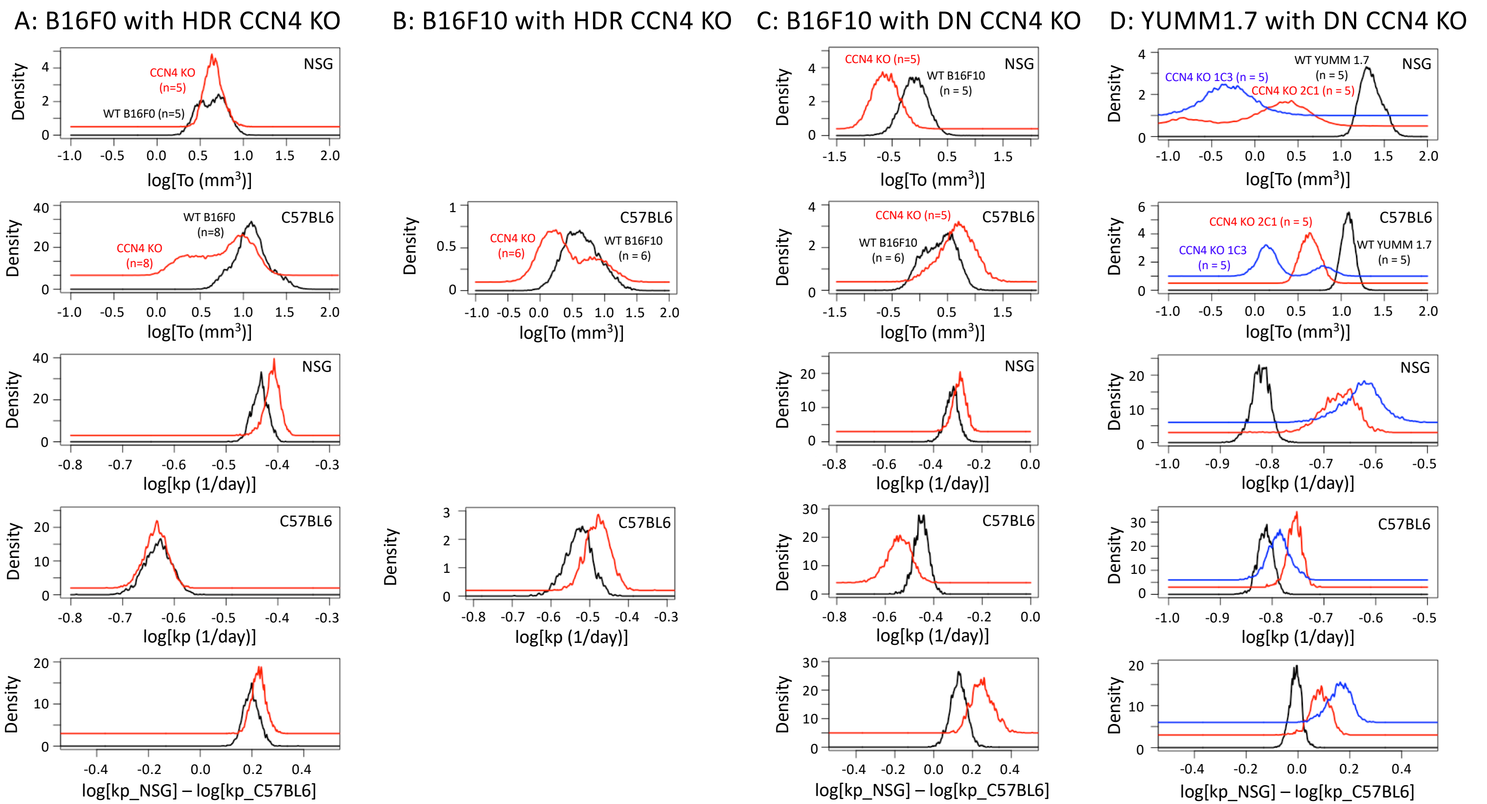


**Supplementary Figure S9. Independent of cell line and CRISPR approach, *Ccn4* knockout consistently reduced the tumor initiation size and increased the growth rate constant, especially in immunocompromised mice.** Comparison of the posterior distributions in the tumor initiation size (To, rows 1 and 2) and growth rate constant (kp, rows 3 and 4) for NSG and C57BL/6crl mice receiving B16F0 wt and HDR Ccn4 KO cells (panel A), B16F10 wt and HDR Ccn4 KO cells (panel B), B16F10 wt and DN Ccn4 KO cells (panel C), and YUMM1.7 wt and DN Ccn4 KO cells (panel D). The difference in growth rate constants for the same cell line variant injected into NSG versus C57BL/6 are shown in row 5, where a positive value indicates that the specific cell line grows faster in NSG than C57BL/6 mice.
